# The Hill function is the universal Hopfield barrier for sharpness of input-output responses

**DOI:** 10.1101/2024.03.27.587054

**Authors:** Rosa Martinez-Corral, Kee-Myoung Nam, Angela H. DePace, Jeremy Gunawardena

## Abstract

The Hill functions, ℋ_*h*_(*x*) = *x*^*h*^*/*(1 + *x*^*h*^), have been widely used in biology for over a century but, with the exception of ℋ_1_, they have had no justification other than as a convenient fit to empirical data. Here, we show that they are the universal limit for the sharpness of any input-output response arising from a Markov process model at thermodynamic equilibrium. Models may represent arbitrary molecular complexity, with multiple ligands, internal states, conformations, co-regulators, etc, under core assumptions that are detailed in the paper. The model output may be any linear combination of steady-state probabilities, with components other than the chosen input ligand held constant. This formulation generalises most of the responses in the literature. We use a coarse-graining method in the graph-theoretic linear framework to show that two sharpness measures for input-output responses fall within an effectively bounded region of the positive quadrant, Ω_*m*_ ⊂ (ℝ^+^)^2^, for any equilibrium model with *m* input binding sites. Ω_*m*_ exhibits a cusp which approaches, but never exceeds, the sharpness of ℋ_*m*_ but the region and the cusp can be exceeded when models are taken away from thermodynamic equilibrium. Such fundamental thermodynamic limits are called Hopfield barriers and our results provide a biophysical justification for the Hill functions as the universal Hopfield barriers for sharpness. Our results also introduce an object, Ω_*m*_, whose structure may be of mathematical interest, and suggest the importance of characterising Hopfield barriers for other forms of cellular information processing.

## 1 Introduction

The Hopfield barrier for an information processing task is the fundamental upper bound to how well that task can be implemented by a mechanism that operates at thermodynamic equilibrium [1]. The only way to exceed this barrier is by expending energy to maintain a steady state away from thermodynamic equilibrium. The existence of such barriers was first pointed out by John Hopfield in his pioneering work on kinetic proofreading for reducing errors in biosynthetic processes like DNA replication [2]. The broader principle just outlined, applicable to any information processing task, is named in his honour [1].

In the present paper, we determine the universal Hopfield barrier for the sharpness of steady-state inputoutput responses. Such responses have been widely used in biochemistry, molecular biology, physiology and pharmacology to quantitatively describe the functional behaviour of biological systems, such as receptors, ion channels, enzymes, transporters, allosteric systems, signalling pathways, gene-regulatory systems, tissues, etc, which interact with an input ligand to produce some output behaviour; see Table S1 in the Supplementary Information (SI). Sharpness, or ultrasensitivity, refers to the amount of output change for a given change in the input, and is often measured by reference to the family of Hill functions,

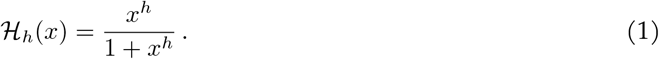

Here, *h >* 0 is the Hill coefficient and *x* is the normalised concentration of the input ligand. The Hill coefficient is frequently quoted as a measure of sharpness, with positive sharpness corresponding to *h >* 1 and negative sharpness to *h <* 1. When *h* = 1, ℋ_1_(*x*) is the classical Michaelis-Menten inputoutput response [3], which is the baseline for the absence of sharpness. Hill functions are also frequently used to represent input-output responses in dynamical system models, where their sharpness underlies the emergence of multiple steady states or limit cycle oscillations [4, 5].

Because Hill functions are so widely used, it is sometimes forgotten that they have no justification when *h* ≠1 [6, 7]. Unlike the Michaelis-Menten response ℋ_1_(*x*), which was originally derived to explain enzyme kinetics and has been found in many other contexts [8], the Hill functions are merely a convenient family of rational functions which Archibald Vivian Hill selected to fit data on the oxygen-binding response of haemoglobin [9]. One of the main results of this paper is to give, for the first time, a rigorous biophysical justification for the Hill functions.

We provide an overview of our approach and results here before explaining the technical details below. We specify input-output responses using the *linear framework*, an approach to Markov processes based on directed graphs with labelled edges [11, 12]; for up-to-date reviews, see [13, 14]. In this approach, graph vertices represent molecular states, directed edges represent transitions and edge labels represent transition rates. Fig.1A shows a graph with a *hypercube structure*, denoted *C*_2+1_, which represents the binding and unbinding of 2 ligands to 3 sites on a biomolecule. We use *structure* to refer to vertices and edges only, disregarding labels. Hypercube-structure graphs frequently underlie models of inputoutput responses [15, 16, 17, 18], including several in Table S1. Edge labels can include terms, such as concentrations of binding ligands, that describe the interaction between the graph and its environment (Fig.1A). The core assumptions about ligands are detailed below.

**Figure 1:**
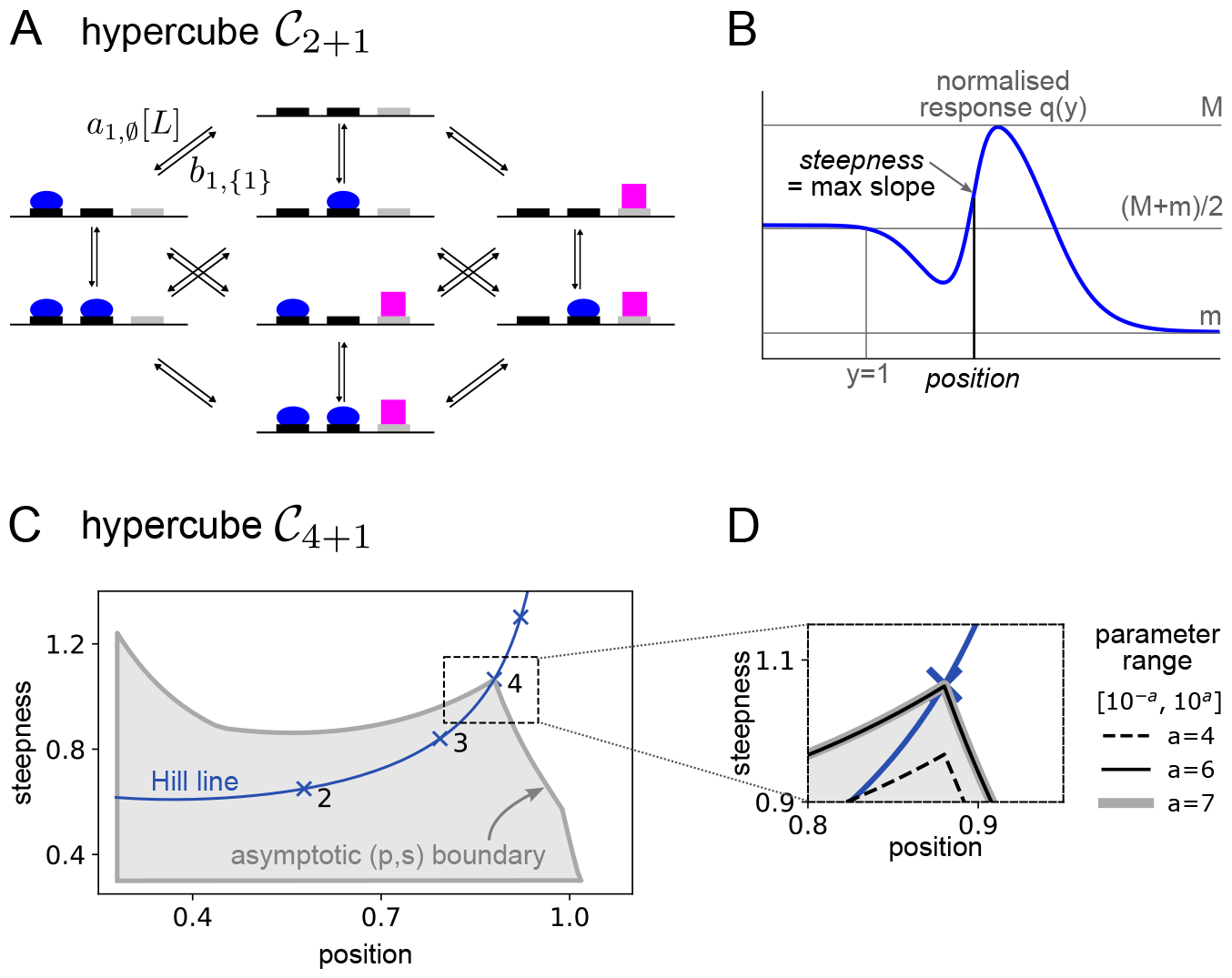
Hypercube-structure graph, definition of position and steepness and position-steepness region. **A**. Linear framework graph with the hypercube structure, *C*_2+1_, representing the binding to a biomolecule of one ligand (*L*, blue oval) to two sites and a second ligand (magenta square) to a third site. Only two edge labels are shown for clarity, with the binding edge label containing a term, [*L*], for the free concentration of ligand *L*. This graph could describe a gene regulation system in which *L* is a transcription factor that recruits RNA Polymerase to a promoter [1, 10]. **B**. Normalised input-output response *q*(*y*) (blue curve), where the normalisation procedure for *y*, depicted in gray font, ensures that *q*(1) = (*M* + *m*)/2 (Eq.10). Steepness is defined as the maximal unsigned slope of *q*(*y*), and position as the smallest value of *y* for which the steepness is attained (Eq.11). **C**. Asymptotic (*p, s*) region for the hypercube *C*_4+1_, obtained by random sampling of parameters over increasing ranges until the boundary of the occupied region stabilises (see inset in panel **D**). The input ligand binds at four sites, the output is the steady-state probability of the fifth site being occupied by its ligand, and the graph is at thermodynamic equilibrium. The region has been truncated to the left and below to focus on the area of interest around the *Hill line* (blue), which is the locus of (*p, s*) points for the Hill functions (Eq.1), with the integer Hill coefficients marked. **D**. Expanded view of the region in panel **C** in the vicinity of the cusp, showing the asymptotic stabilisation as the parametric range increases (see also Figs.S4, S5).

The linear framework allows the steady-state probabilities of graph vertices to be calculated as rational algebraic functions of the labels (Eq.4). Importantly, this can be done whether or not the steady state is one of thermodynamic equilibrium, a property that is determined by the edge labels. An output response can then be defined as a non-negative linear combination of the steady-state probabilities and considered as a function of the concentration of a chosen input ligand, with all other concentrations kept constant (Eq.5).

In previous work, we introduced two intrinsic, non-dimensional measures, *position* and *steepness*, to quantify the sharpness of such an input-output response [1], as described in Fig.1B. By sampling parameter values appropriately, we plotted the two-dimensional position-steepness, or (*p, s*), regions for various input-output responses on hypercube-structure graphs with different numbers of input binding sites, assuming the corresponding systems were at thermodynamic equilibrium [1, 10]. Fig.1C shows part of the (*p, s*) region for a hypercube-structure graph similar to that in Fig.1A but with 4 binding sites for the input ligand.

We found that these (*p, s*) regions exhibit four characteristic properties. First, the regions have an asymptotic boundary: if the range over which the model parameters are sampled is steadily increased, the boundary of the (*p, s*) region stabilises (Fig.1D), giving rise to the asymptotic boundary and the asymptotic region (Fig.1C). Second, the boundary encloses a region that is *effectively bounded* in the positive quadrant. The (*p, s*) region is not bounded in the whole positive quadrant ℝ^+^ *×* ℝ^+^; there are “wings” that become asymptotic to the axes. These wings are not shown in Fig.1C; they are not the focus of this paper and appear not to be biologically relevant. However, given *a >* 0, no matter how small, that part of the (*p, s*) region which falls within [*a*, ∞) *×* [*a*, ∞) is bounded, which is what we mean by “effectively bounded”. Third, these (*p, s*) regions exhibit a cusp that falls on the *Hill line*, the locus of (*p, s*) points for the Hill functions. The tip of the cusp lies below the (*p, s*) point with Hill coefficient equal to the number of input binding sites and approaches this Hill point more closely as the parametric range increases (Fig.1D). In other words, if the system has *m* binding sites for the input ligand, the (*p, s*) point for ℋ_*m*_ acts as a barrier to sharpness. Note the importance of using two sharpness measures to draw this conclusion: each one of the measures can individually exceed the corresponding value for ℋ_*m*_ but they cannot both do so simultaneously (Fig.1C). Fourth, parameter values can be found away from thermodynamic equilibrium whose (*p, s*) points lie above and to the right of the (*p, s*) point of ℋ_*m*_. This confirms that ℋ_*m*_ is the Hopfield barrier for sharpness of input-output responses on hypercube-structure graphs like those in Fig.1A.

Results of this kind already constrain experimental data. If the data fall outside the (*p, s*) region, then no model of this kind can account for the data, no matter what parameter values are chosen, and this can be asserted without fitting the model to the data [10]. However, the scope of this conclusion is limited by the underlying hypercube structure. Much greater molecular complexity may actually be present, such as co-regulators, conformations, internal states, modifications, etc (Fig. S1). A different model will typically yield a different (*p, s*) region (Fig. S6) and this region may be able to account for the data.

We will show here that this limitation may be overcome for sharpness at thermodynamic equilibrium. We use a method of coarse graining in the linear framework to show that there is a *universal* bounded region that contains the (*p, s*) point for the input-output response of any Markov process model at thermodynamic equilibrium and this region exhibits the same four properties described above. The model may be arbitrarily complicated, subject to the core assumptions detailed below. In particular, the Hill function emerges as the universal, model-independent Hopfield barrier for sharpness.

Most studies in the biological literature focus on specific models and it is rare to be able to make a rigorous claim about all models within a large and widely-used class, such as the Markov process models studied here. The universality that we have uncovered for sharpness may potentially hold more widely and suggests new directions to explore in the mathematics and biophysics of cellular information processing (Discussion).

## 2 Results

### 2.1 Graphs, Markov processes and input-output responses

The linear framework was introduced in [11, 12] and reviewed in [19, 13, 14]; the Methods and the SI provide more details. Under the *reservoir assumptions* defined below, linear framework graphs are equivalent to finite-state, continuous-time, time-homogeneous Markov processes that have infinitesimal generators [12, Theorem 4]. The graph specifies the master equation of the Markov process (Methods, Eq.13), which is a linear differential equation from which the framework acquires its name. Graph vertices represent the states of the Markov process. There is an edge between two vertices when the infinitesimal rate for this transition is positive, in which case this positive rate becomes the edge label, with dimensions of (time)^−1^. Vertices are typically denoted by 1, *· · ·, n*, edges by *i* → *j* and labels by ℓ(*i* → *j*). Edge labels can contain terms that describe the interaction between the graph and its environment (Fig.1A). Fig. S1 shows some of the molecular complexity that may be accommodated within the graph formalism. From now on, we will refer interchangeably to graphs and their corresponding Markov processes.

The use of graphs to study Markov processes has its roots in the pioneering work of Hill [20] and Schnakenberg [21]. It is rarely seen in the Markov process literature and has only occasionally appeared in the biophysics literature [22], until the development of the linear framework [23, 24, 25, 26]. The main distinction in the linear framework approach is to treat the graph as a mathematical object in its own right, in terms of which results can be formulated, which, as we will see here, can accommodate some of the molecular complexity found in biology.

We use “ligand” to refer to any component in the environment that interacts with the graph through binding and unbinding, like those represented by the blue oval and magenta square in Fig.1A. Depending on the context, such as gene regulation, a ligand may be a transcription factor, an enzyme complex like RNA Polymerase, a co-regulator like Mediator, a nucleosome, etc [27]. Ligand binding is assumed to follow mass action and to be first order, so that a binding edge label acquires a term for the free ligand concentration (Fig.1A). Ligands are assumed not to engage in activities outside the graph, such as oligomerisation; such activities may be accommodated [28] but complicate the arguments given here. Most importantly, ligands are assumed to be present in sufficient quantity that binding does not appreciably change their free concentration. This *reservoir assumption*, which is implicitly made in all treatments of input-output responses, is similar to the assumption in classical thermodynamics of a heat bath, with which energy can be exchanged without altering the temperature. First-order binding and reservoirs are the core assumptions that underlie all the models and results of this paper; they are commonly used in the literature, not always explicitly.

The linear framework enables the steady-state (s.s.) probability, 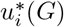, of vertex *i* of graph *G* to be calculated as a rational function of the edge labels. Recall that *G* is *strongly connected* if any two distinct vertices, *i* ≠*j*, are connected by a directed path, *i* = *i*_1_ → *i*_2_ → *· · ·* → *i*_*k*_ = *j*. Provided *G* is strongly connected, there is a unique s.s., which is described up to a proportionality constant by the vector, *ρ*(*G*), with components

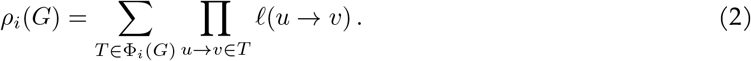

Here, F_*i*_(*G*) is the set of *spanning trees* of *G* that are *rooted* at *i*. A spanning tree is a subgraph of *G* that includes every vertex (spanning), has no cycles when edge directions are ignored (tree) and has only one vertex with no outgoing edge (the root). The s.s. probability is recovered from Eq.2 by normalising, as in Eq.4 below.

Eq.2 shows that s.s. probabilities depend on all the edge labels in the graph and are subject to a combinatorial explosion even for relatively small graphs, which arises from having to enumerate all spanning trees: the structure 𝒞_4_, for example, has 42, 467, 328 spanning trees rooted at each vertex [29]. However, a substantial simplification occurs if *G* can reach a s.s. of thermodynamic equilibrium (t.e.). A graph *G* is at t.e. if two conditions are satisfied. First, *G* is *reversible*, so that if *i* → *j*, then the reverse transition, *j* → *i*, is also present. Second, *detailed balance* holds, so that any pair of reversible edges, *i* ⇋ *j*, is independently in flux balance: 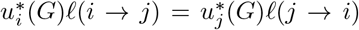. Reaching a s.s. in which detailed balance holds is equivalent to the following *cycle condition* on the labels of a reversible graph. Let *P* be any path of reversible edges, *P* : *i*_1_ ⇋ *i*_2_ ⇋ *· · ·* ⇋ *i*_*k*_, and let *µ*(*P*) denote the product of the label ratios along *P*,

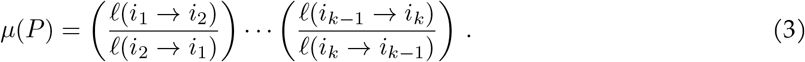

The cycle condition requires that *µ*(*P*) = 1 whenever the path is a cycle, with *i*_*k*_ = *i*_1_. The quantity log *µ*(*P*) is interpreted in stochastic thermodynamics as the *entropy* generated along *P* [30], so that t.e. corresponds to there being no entropy generation over cycles in *G*.

At t.e., an alternative vector, *µ*(*G*), may be used to calculate steady state probabilities. Choose a reference vertex, which we will index as 1. In principle, this can be any vertex but it will be convenient to choose one in which no input binding site is bound (Methods). Now choose any path, *P*_*i*_, of reversible edges from 1 to *i*; and let *µ*_*i*_(*G*) = *µ*(*P*_*i*_). The cycle condition ensures that this is well defined. The s.s. probability can then be determined by normalising,

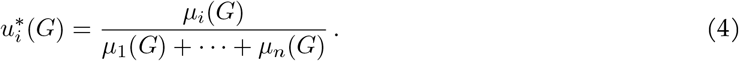

This normalisation can be done either with *µ*(*G*) at t.e., as shown in Eq.4, or with *ρ*(*G*) in the general case. Either way, the resulting expression is a rational function of the edge labels.

When *G* can reach t.e., the quantities log *µ*_*i*_(*G*) can be interpreted in terms of the free energy of vertex *i* relative to the reference vertex 1. Eq.4 then recovers the classical formula of equilibrium statistical mechanics: the denominator is the partition function for the grand canonical ensemble and the terms *µ*_*i*_(*G*) provide the Boltzmann factors. A key advantage of the linear framework is that it reduces to equilibrium statistical mechanics at t.e. but it also enables s.s. probabilities to be exactly calculated away from t.e. by using Eq.2.

Input-output responses on *G* may now be defined by choosing some ligand as input. We denote its concentration by *x*. We assume that *x* is changed quasi-statically—in small increments and sufficiently slowly that the graph relaxes back to a steady state after each change—which fits the conditions under which input-output responses have been measured (Table S1). The concentrations of any other ligands are assumed to be held constant. The output can be any non-negative linear combination of s.s. probabilities, considered as a function of *x*,

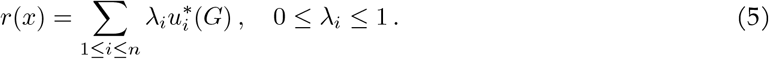

The restrictions ensure that *r*(*x*) is non-dimensional and normalised to lie in [0, 1]. It follows from Eqs.2 and 4 that *r*(*x*) is a rational function of *x*.

### 2.2 Coarse graining

In this section, we will show that if *G* is any strongly connected, reversible graph that reaches t.e., and *r*(*x*) is any input-output response on *G*, then *r*(*x*) can be rewritten as an input-output response on some reversible substructure of the hypercube 𝒞_*m*_, where *m* is the number of input binding sites, with edge labels that satisfy the cycle condition. This result begins to explain how universality arises at t.e.: no matter how complex the input-output response, it is mathematically equivalent to one that involves only the binding and unbinding of the input ligand. This rewriting requires finding edge labels for the substructure of 𝒞_*m*_, as well as the appropriate coefficients for its input-output response. The coarse-graining strategy introduced in [31] provides the necessary approach. We apply it here with further details in the Methods. To avoid trivial special cases, we assume from now on that *m >* 1.

Let *L* be the input ligand and let ν (*G*) denote the set of vertices of *G*. Coarse graining does not require *G* to satisfy the cycle condition, although we will make this assumption later. Coarse graining starts from any partition of the vertices of *G* into disjoint subsets and constructs a linear framework graph, *C*(*G*), whose vertices are the subsets of the partition. We choose the partition given by collecting together those vertices with the same pattern of binding of *L*. Binding patterns are indexed by subsets *S* ⊆ {1, *· · ·, m*}. Let *G*_*S*_ ⊆ *ν* (*G*) contain those vertices *i* ∈ *ν* (*G*) such that, if *s* ∉ *S* then *L* is bound at *s* in vertex *i* but if *s ϵ* ∈ *S* then *L* is not bound at *s* in vertex *i*. The vertex *i* may have many other features as a vertex of *G* in addition to the sites bound by *L* but this coarse graining ignores them. There is an edge in *C*(*G*), *w* →_*C*(*G*)_ *z* if, and only if, there is an edge in *G, i* →_*G*_ *j*, for some vertex *i* ∈ *G*_*w*_ and some vertex *j* ∈ *G*_*z*_. It follows that the vertices and edges of *C*(*G*) are those of the hypercube structure *C*_*m*_. But *C*(*G*) may not be all of *C*_*m*_. This can happen because of mutual exclusion (Fig. S1B), in which some vertices of *C*_*m*_ are not reached, or because of ordering (Fig. S1C), in which some edges of 𝒞_*m*_ are not used. Accordingly, the structure of *C*(*G*) is generally only a substructure of 𝒞_*m*_. Fig.2 shows an example of coarse graining in which *C*(*G*) is all of 𝒞_2_.

**Figure 2:**
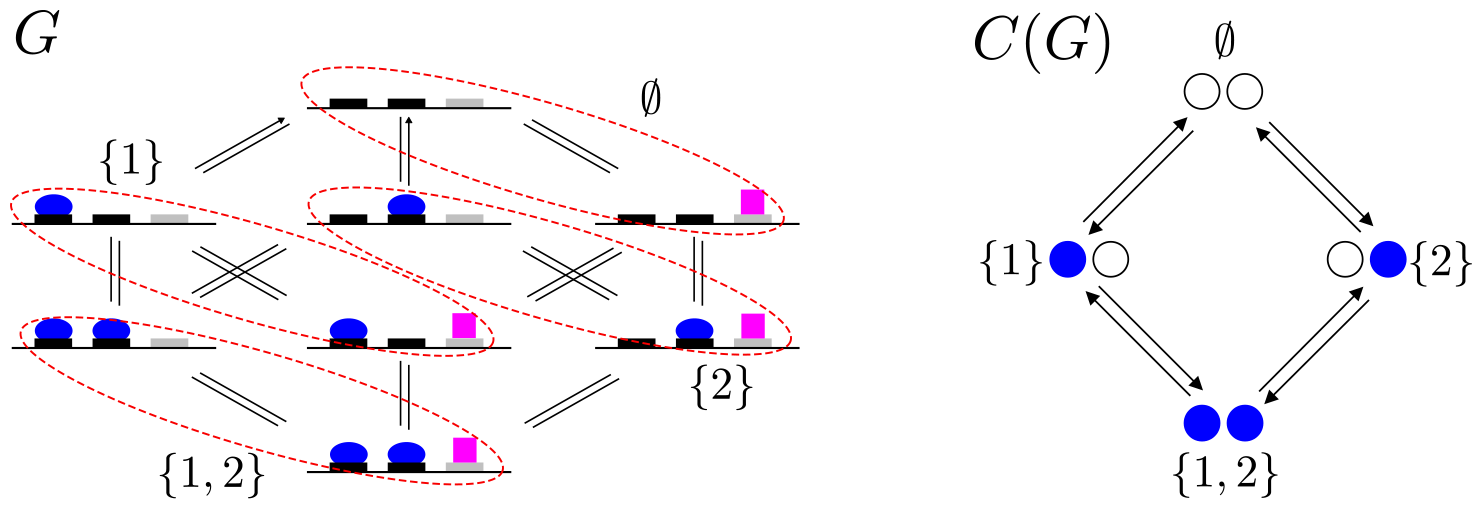
Coarse graining, showing only graph structures. On the left, the structure *G*, from Fig.1A, is being coarse grained, as described in the text, with the blue oval as the input ligand *L*. The vertices of *G* are partitioned into subsets (red, dashed ovals) corresponding to the patterns of binding of *L* to *m* = 2 sites; the magenta square is ignored. The input binding sites are indexed 1 and 2 from left to right and the subsets of sites are indexed in set notation ∅, {1}, {2}, {1, 2}. The resulting coarse-grained structure, *C*(*G*), is shown on the right, with vertices indexed by the corresponding subsets. In this case the full structure of the hypercube 𝒞_2_ is recovered. The edges and labels of *C*(*G*) are explained in the text and the SI.

It can be shown that labels may be assigned to the edges of *C*(*G*) in essentially only one way (Methods, Eq.14), such that *C*(*G*) satisfies the cycle condition and the following coarse-graining equation holds [31],

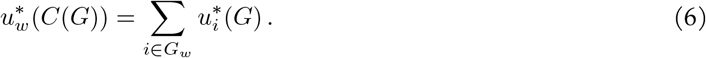

Eq.6 is what would be expected from a coarse-graining at s.s. Eq.5 may now be rewritten as,

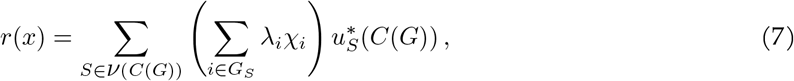

where χ_*i*_ can be seen from Eq.6 as the s.s. probability of *i* conditioned on the subset *G*_*S*_ ⊆ *ν* (*G*),

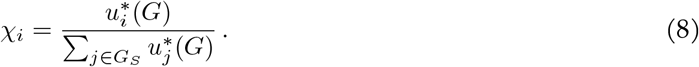

The terms χ_*i*_ may be extremely complicated in general, as they summarise the other features that are present in *G*. The key point, however, is that if *G* does satisfy the cycle condition, then the χ_*i*_ do not depend on *x* (Methods). It follows that, provided that *G* satisfies the cycle condition, Eq.7 expresses *r*(*x*) as a valid input-output response on a graph that is a substructure of 𝒞_*m*_, as claimed above. Let us call this graph 𝒢_*m*_.

A universal (*p, s*) region can now be generated by sampling not only the labels of 𝒢_*m*_ but also the coefficients *λ*_*i*_ (Eq.5) that appear in the input-output responses on 𝒢_*m*_. However, a further simplification arises because *r*(*x*) has the following rational structure (Methods),

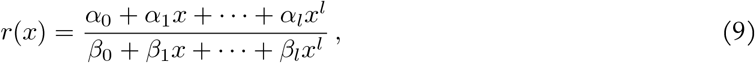

where *l* is the maximum number of sites bound by *L*, so that 1 ≤ *l* ≤ *m*; the denominator coefficients are all positive, 0 *< β*_*i*_; and the numerator coefficients are non-negative and not greater than the corresponding coefficients in the denominator, 0 ≤ *α*_*i*_ ≤ *β*_*i*_. It can be shown that any choice of *α*_*i*_ and *β*_*i*_ that satisfies these conditions corresponds to an equilibrium input-output response (SI), so that Eq.9 exactly describes the equilibrium input-output responses of Markov process systems with *m* input binding sites.

Eq.9 shows that the rational structure of an equilibrium input-output response is largely independent of the graph *G* from which it is derived. *G* determines the coefficients, *α*_*i*_, *β*_*i*_, but the degree in *x* of the denominator of *r*(*x*), namely *l*, depends only on the maximum number of sites bound by the input, irrespective of the complexity of *G*. The rational structure of Eq.9 is a preliminary mathematical expression of the universal Hopfield barrier and is the basis for analysing sharpness below.

The non-dependence of χ_*i*_ on *x*, which is crucial for the structure of *r*(*x*) described in Eq.9, breaks down emphatically if *G* does not satisfy the cycle condition. The resulting *r*(*x*) can then no longer be an inputoutput response on some substructure of 𝒞_*m*_. The algebraic structure of non-equilibrium input-output responses is strikingly different, as we will see below.

### 2.3 Intrinsic measures of sharpness

To define measures of sharpness, it is necessary to normalise the input-output response. The output value is normalised already in the light of Eq.5. Since there is no naturally independent quantity against which to normalise the input concentration, *x*, its normalisation has to be intrinsically determined for each response. The input-output responses allowed by Eq.9 can be non-monotonic and complicated (Fig.1B).

Accordingly, we choose the normalisation value, denoted *x*_0.5_, to be the smallest positive value of *x* at which the response is halfway between its supremum and its infimum, which exist because 0 ≤ *r*(*x*) ≤ 1. More precisely, we define (Figs.1B and S2),

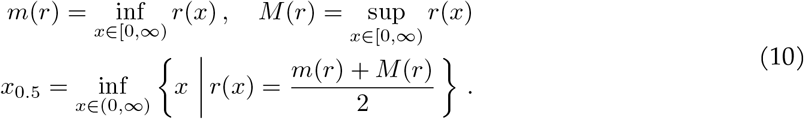

We explicitly choose *x*_0.5_ *>* 0. This can always be done because, even if *r*(0) = (*m*(*r*) + *M* (*r*))/2, there must be *x >* 0 for which *r*(*x*) has the same value. The normalised response, *q*(*y*), where *y* = *x/x*_0.5_, is then defined by *q*(*y*) = *r*(*yx*_0.5_). Note that *x*_0.5_ depends on *r*.

Following normalisation, the two intrinsic measures of sharpness are the supremum of the absolute value of the derivative of *q*(*y*), which we call *steepness* and denote *s*(*r*), and the smallest *y* value that attains the supremum, which we call *position* and denote *p*(*r*). The supremum is attained at a finite value of *y* (SI), so that,

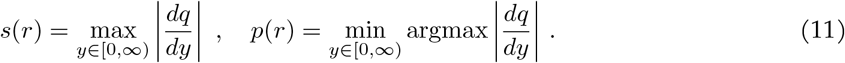

Because of the dependence of *x*_0.5_ on *r*, position and steepness are scale invariant: *p*(*r*(*cx*)) = *p*(*r*) and *s*(*r*(*cx*)) = *s*(*r*), for any scale factor *c >* 0. If we denote by *s*^*u*^(*r*) and *p*^*u*^(*r*) the un-normalised versions of steepness and position, obtained using *dr/dx* in place of *dq/dy* in Eq.11, then the relationship between the normalised and un-normalised versions is given by a complementary scaling by *x*_0.5_: the steepness is multiplied and the position is divided,

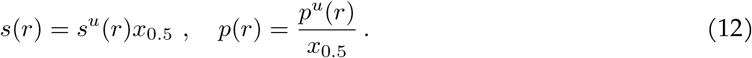

This relationship will be helpful to interpret (*p, s*) regions below.

### 2.4 Universal position-steepness region and the Hopfield barrier

We elaborated the techniques previously introduced to estimate (*p, s*) regions like that in Fig.1C [1, 10, 32] to plot the universal (*p, s*) region, Ω_*m*_, for input-output responses at t.e. with *m* input binding sites (Fig.3A). The coefficients in Eq.9 were sampled for *l* = *m*; the (*p, s*) points of the corresponding rational functions were plotted; and the resulting region was grown by biasing the sampling and expanding the parametric range so as to establish the asymptotic boundary. The algorithm is summarised in the Methods with further details in the SI.

Fig.3A shows Ω_4_, with confirmation of the asymptotic boundary shown in Fig. S4, and Fig. S5 shows the same for Ω_6_. These universal regions have the same characteristic properties satisfied by the *C*_4+1_ model in Fig.1C, as described in the Introduction. First, the regions have an asymptotic boundary. Second, the regions are effectively bounded in the positive quadrant. The wings that asymptote to the axes are more visible in Fig.3A. Third, Ω_*m*_ has a cusp that falls on the Hill line and lies just below the (*p, s*) point of *H*_*m*_. During asymptotic convergence, the cusp approaches increasingly close to the (*p, s*) point of *H*_*m*_ as the parametric range is increased (Figs. S4 and S5). As before, the Hill function acts as a sharpness barrier at t.e.: while there are input-output responses with either higher position or higher steepness than *H*_*m*_, there are none with both higher position and higher steepness.

The coefficient constraints for Eq.9, which characterise equilibrium input-output responses (SI), are required for these properties of Ω_*m*_. If rational functions are permitted which do not obey the constraints, then (*p, s*) points can be found that lie outside the universal region (Fig.3B). This confirms that the asymptotic region arises only for equilibrium input-output responses. As a further check on the universality of Ω_*m*_, we calculated the (*p, s*) regions for six specific models with *m* = 4 input binding sites and found them all to be contained within Ω_4_, as expected (Fig. S6). Details of the models are given in the SI.

The situation is profoundly different away from t.e., as discussed in the Introduction. We considered the graph with hypercube structure *C*_4_ and the input-output response given by fractional saturation, which is the average number of bound inputs normalised to the total number of input binding sites (here 4). When edge labels are allowed to be away from t.e., we readily found (*p, s*) points that lie outside Ω_4_ (Fig.3C). In particular, we found (*p, s*) points that are greater in both position and steepness than those of *H*_4_ (Fig.3C, red points), thereby confirming that ℋ_4_ is the universal Hopfield barrier for sharpness of input-output responses with *m* = 4 input binding sites. There is nothing special about the structure *C*_4_: other graph structures with *m* = 4, such as those in Fig. S6, can yield (*p, s*) points lying outside Ω_4_ and exceeding ℋ_4_ in both position and steepness, when edge labels are allowed to be away from t.e.

The tapering wings of Ω_*m*_ (Fig.3A) can be understood as follows. Recall the un-normalised versions of steepness and position, *s*^*u*^(*r*) and *p*^*u*^(*r*), which satisfy Eq.12. We can distinguish two extreme cases, when *p*^*u*^(*r*) ≪ *x*_0.5_, so that *p*(*r*) = *p*^*u*^(*r*)*/x*_0.5_ is small, while *s*(*r*) = *s*^*u*^(*r*)*x*_0.5_ can become large, or when *x*_0.5_ ≪ *p*^*u*^(*r*), so that *p*(*r*) is large, while *s*(*r*) can become small. Input-output responses that satisfy these conditions appear not to be biologically meaningful.

### 2.5 Failure of universality away from thermodynamic equilibrium

As discussed above, the universality of Ω_*m*_ breaks down completely away from t.e. This strikingly different behaviour may be understood in terms of the difference between the vectors *µ*(*G*) at t.e. (defined through Eq.3) and *ρ*(*G*) away from t.e. (Eq.2), in terms of which s.s. probabilities are calculated by normalising (Eq.4). For a given vertex *i* ∈ *ν* (*G*), *µ*_*i*_(*G*) = *µ*(*P*), where *P* is a path from 1 to *i*, with the cycle condition ensuring that this value is independent of the chosen path. It is this property of *path independence* that ultimately shows, through coarse graining, that the degree of *x* in *µ*_*i*_(*G*) is given simply by the number of input binding sites that are bound by the input in vertex *i* (Methods). This degree is essentially independent of the structure of the graph *G*. It then readily follows that the rational structure of input-output responses at t.e., as described in Eq.9, is also essentially independent of *G*.

Away from t.e., however, we must use *ρ*(*G*), rather than *µ*(*G*), to calculate s.s. probabilities. An inputoutput response is still a rational function of *x* but, as Eq.2 makes clear, *ρ*_*i*_(*G*) depends on all spanning trees rooted at *i*. In consequence, the degree of *x* in *ρ*_*i*_(*G*) now depends crucially on the structure of *G* and becomes unrelated to the number of bound sites in *i* (SI). For hypercube graphs of structure *C*_*m*_, inputoutput responses away from t.e. have degree 2^*m*^ − 1 in *x* [1] and can therefore have substantially higher sharpness than responses at t.e. [25]. It is this marked difference in rational structure that underlies Fig.3C and explains the failure of universality away from t.e.

Very little is known about the shape of (*p, s*) regions away from t.e., which, as just explained, are now model dependent. Numerical estimation of regions beyond quite small graphs is hampered in part by the combinatorial intractability of Eq.2. What little evidence there is [1] suggests that non-equilibrium (*p, s*) regions also have a cusp on the Hill line, just below the (*p, s*) point for ℋ_*z*_ where *z* is an integer.

However, it is an open problem to understand how *z* depends on the underlying graph.

## 3 Discussion

We have provided here, for the first time, a rigorous biophysical justification for the Hill functions. As pointed out in the Introduction, they have been widely exploited in biology for over a century, for both data fitting and modelling. Yet, they have been nothing other than a convenient family of rational functions. We have shown by numerical calculations for *m* = 4 (Fig.3) and *m* = 6 (SI) that Hill functions with integer Hill coefficients are the universal Hopfield barriers for sharpness of input-output responses: given any Markov process model with *m* input binding sites at t.e., no matter how complicated, the sharpness of any input-output response (Eq.5) lies within the universal, model-independent (*p, s*) region Ω_*m*_, and cannot be higher in both position and steepness than that of the Hill function ℋ_*m*_ (Fig.3A). In contrast, if any such graph is away from t.e., then input-output responses can be found whose position and steepness both exceed those of ℋ_*m*_ (Fig.3C). A. V. Hill could not have anticipated, at the time he introduced his eponymous functions [9], their deep connection to thermodynamics.

Our numerical results strongly suggest that the conclusions described above hold for all values of *m* and it remains an open problem to give a mathematical proof of this. Considerable subtlety arises because of the shape of Ω_*m*_. It is not true in general that the position or steepness of an input-output response is less than the position or steepness, respectively, of ℋ_*m*_ but both assertions become true within the cusp (Fig.3).

Position and steepness become increasingly tightly constrained within the cusp so as to asymptotically fall on the Hill line itself. The precise nature of this changing constraint is not yet understood and this appears to be one of the main barriers to a proof.

The winged, cuspidal shape of Ω_*m*_ (Fig.3A) is particularly tantalising. Its universality suggests that it may have some deeper mathematical significance that has yet to be understood. Perhaps this may encourage mathematicians to examine more closely a mathematical object that has emerged directly from biology. There remains much work to be done, as noted above, to understand the sharpness regions of input-output responses away from t.e. Another important question arises in moving beyond the reservoir assumptions made here. Biological ligands are always present in limited amounts and may be engaged in other activities beyond the system of interest. Such issues have been largely ignored in the literature but evidence is emerging as to the consequences of doing so [33, 34]. Ligand limitation and distraction can be accommodated within the linear framework [28] but the resulting input-output responses begin to stray outside the elegant confines of rational functions.

Our results illustrate the significance of Hopfield’s insights into energy expenditure, as first put forward for biosynthetic error correction [2] and then elaborated, as explained in the Introduction, for any form of information processing [1]. No matter what information processing task is being undertaken, there is a fundamental limit—the Hopfield barrier—to how well it can be carried out at t.e. The limit is set by fundamental physics, in effect by the cycle condition. Energy expenditure has been widely studied in areas like pattern formation, force generation and active matter [35, 36] but its role in information processing has been more elusive. This may reflect the fact that, in areas other than information processing, the relevant Hopfield barriers are zero: for example, directed movement is impossible at t.e. Information processing, in contrast, can certainly occur at t.e., even though, as Hopfield recognised, evolution has bypassed the Hopfield barriers.

We believe the time is now ripe to analyse in more depth the functional impact of energy expenditure in cellular information processing. Previous studies have suggested putative Hopfield barriers [37, 38, 39, 23] and there is now growing evidence for the significance of non-equilibrium functionality in gene regulation [28, 10, 40, 41]. Much insight could be gained by characterising the Hopfield barriers for the various information processing tasks undertaken by cells, as we have done here for the sharpness of input-output responses. Such a research programme may not only bring to light some of the general principles at work in biology but may also reveal further objects of mathematical interest. Moreover, the method of coarse graining used here, which is generally applicable, leads us to ask whether similar universality and model independence may also be found for other Hopfield barriers. We hope the results of the present paper will stimulate further studies of Hopfield barriers in cellular information processing.

## Acknowledgements

We thank Isabel Thomas and Felix Wong for comments on non-equilibrium (*p, s*) regions; an anonymous reviewer for suggestions that improved the paper’s clarity; and Harvard Medical School’s Research Computing Group for access to the O2 High Performance Compute Cluster. RM-C, K-MN, AHD and JG were supported by US National Institutes of Health (NIH) award GM122928 and RM-C was also supported by EMBO Fellowship ALTF683-2019. RM-C acknowledges support of the Spanish Ministry of Science and Innovation through the Centro de Excelencia Severo Ochoa (CEX2020-001049-S, MCIN/AEI /10.13039/501100011033), the Generalitat de Catalunya through the CERCA programme, and grant RYC2021-033860-I funded by MCIN/AEI/10.13039/501100011033 and by European Union NextGenerationEU/PRTR. AHD is a Scientific Program Officer of the Howard Hughes Medical Institute.

## 4 Methods

### 4.1 Master equation and Eq.2

A linear framework graph, *G*, gives rise to a linear dynamics as follows: each edge may be thought of as a chemical reaction under mass-action kinetics with the edge label as the rate constant. Since an edge has only a single source vertex, the dynamics must be linear. It may be written in matrix form as

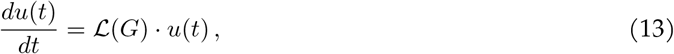

where *u*(*t*) is the vector of vertex probabilities at time *t* and ℒ (*G*) is the Laplacian matrix of *G* [42]. Under reservoir assumptions, Eq.13 is the master equation of the corresponding Markov process [12, Theorem 4]. A s.s. of Eq.13 must lie in the kernel of ℒ (*G*), which is 1-dimensional when *G* is strongly connected. The canonical basis element *ρ*(*G*) ∈ ker ℒ (*G*) is calculated by using the Matrix-Tree theorem of graph theory, which relates the minors of ℒ (*G*) to spanning trees of *G*. This gives Eq.2, from which the s.s. can be calculated by normalising to remove the proportionality constant, as in Eq.4.

### 4.2 Coarse-graining

This method was introduced in [31]. Let *G* be any strongly connected, reversible graph. Choose any partition of the vertices into disjoint subsets: ν (*G*) = *G*_1_ ∪ *· · ·* ∪ *G*_*s*_ and *G*_*w*_ ∩ *G*_*z*_ = ∅ when *w ≠z*. A *coarse-grained* graph, *C*(*G*), is constructed on the vertices 1, *· · ·, s*, corresponding to the subsets of the partition. There is an edge *w* →_*C*(*G*)_ *z* if, and only if, there is an edge *i* →_*G*_ *j* for some vertex *i* ∈ *G*_*w*_ and some vertex *j* ∈ *G*_*z*_. *C*(*G*) thereby inherits reversibility from *G*. The edge labels on *C*(*G*) are given by,

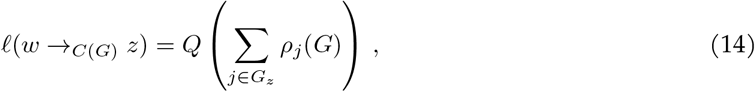

where *ρ*(*G*) is the vector defined in Eq.2. The quantity *Q* in Eq.14 is chosen to ensure that the labels have dimensions of (time)^−1^ but its actual value is irrelevant because, with these labels, *C*(*G*) satisfies the cycle condition, even when *G* does not. Hence, as far as s.s. probabilities of *C*(*G*) are concerned, only the label ratios are relevant (Eq.4), so that *Q* cancels out. The key point is that, with the labelling in Eq.14, the coarse-graining formula in Eq.6 holds. The choice of labels in Eq.14 is essentially unique if *C*(*G*) has to satisfy the cycle condition and Eq.6 has to hold. Note that this coarse graining is only at s.s. and nothing is implied about the dynamics of *C*(*G*).

### 4.3 Rational structure and Eqs.8 and 9

The independence of *χ*_*i*_ in Eq.8 from *x* arises because, if *j* ∈ *G*_*S*_, then 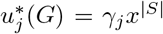, where |*S*| is the size of *S*, or the number of input binding sites that are bound by the input, and *γ*_*j*_ is independent of *x*. To see this, recall that the reference vertex, 1, in *G* was chosen to be a state in which no input ligand is bound. Take any path, *P*, of reversible edges from 1 to *j, P* : 1 = *i*_1_ ⇋ *· · ·* ⇋ *i*_*k*_ = *j*. It follows from the definition of *µ*(*P*) in Eq.3 that traversing *P* from *i*_1_ to *i*_*k*_, forward edges may be encountered at which the input binds, which each contribute a factor *x* to *µ*(*P*), as well as forward edges at which the input unbinds, which each contribute a factor *x*^−1^ to *µ*(*P*). Since no ligand is bound in *i*_1_ = 1 and *j* ∈ *G*_*S*_ has |*S*| input binding sites, the net effect of the bindings and unbindings along *P* must be to contribute exactly *x*^|*S*|^ to *µ*(*P*). Provided *G* satisfies the cycle condition, *µ*(*P*) is independent of the choice of *P*. Hence, 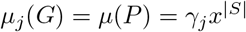, where *γ*_*j*_ does not depend on *x*. It then follows that *x* occurs to the same degree in both the numerator and each term of the the denominator of Eq.8, so that it cancels out and χ_*i*_ is independent of *x*, as claimed.The denominator of 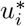 in Eq.4 is now a polynomial in *x* of degree *l*, where *l* is the maximum number of input binding sites that are bound by the input ligand. Also, every degree less than *l* must occur in the denominator, since states are formed by successive binding of the input ligand. Hence, from Eq.5, *r*(*x*) is a rational function whose denominator polynomial is of degree *l* in *x*, as shown in Eq.9, with *β*_*i*_ *>* 0 for 0 ≤ *i* ≤ *l*. Since the numerator of 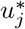 in Eq.4 is always part of the denominator, it follows from Eq.5 that 0 ≤ *α*_*i*_ ≤ *β*_*i*_.

### 4.4 Determination of the universal (*p, s*) region in Fig.3A

Parameters are sampled as follows. Eq.9 has 2(*m* + 1) parameters (coefficients) for graphs at t.e. with *m* input binding sites. The denominator parameters, *β*_*i*_, are sampled by choosing log_10_ *β*_*i*_ uniformly at random in the interval [−*a, a*], for a fixed exponent range *a*. Having chosen the *β*_*i*_, the logarithm of the numerator parameters, log_10_α_*i*_, are sampled uniformly at random in the interval [−*a*, log_10_ *β*_*i*_], to satisfy the constraints in Eq.9. As previously found [10], the boundary of the (*p, s*) region stabilises rapidly as *a* is increased (Fig. S4) to give an asymptotic boundary.

Boundaries are estimated as follows. The 2-dimensional (*p, s*) space is divided into a grid of small square cells of side length 0.005. The current working boundary is defined by those cells which contain sampled (*p, s*) points but which have only empty cells above or below in the same column or to the left or right in the same row. The working boundary is then recomputed in two phases. First, each of the sampled parameter sets that yield (*p, s*) points on the working boundary is repeatedly “mutated” by randomly choosing new parameter values near the sampled value, independently for each parameter, until a parameter set is found whose (*p, s*) point goes into an empty cell. This may generate a new working boundary. Second, for each sampled (*p, s*) point on the resulting boundary, a target point is determined that lies outside the boundary and repeated mutations are attempted, as before, to reduce the distance in (*p, s*) space to this target point. This second phase is important to avoid becoming trapped in deep valleys during the first phase. The algorithm is considered to converge when no new boundary cells are created after a number of iterations that is specified as a hyperparameter; we took it to be 1500.

## Supplementary Information

We follow the notation and terminology of the main text, using s.s. for steady state and t.e. for thermodynamic equilibrium.

## 1 Input-output responses

Table S1 shows example input-output responses from a variety of biological contexts. Similar *dose responses* are also commonly measured in toxicology, where the inputs are diverse chemicals and outputs can be at the organism level [1].

**Table S1:**
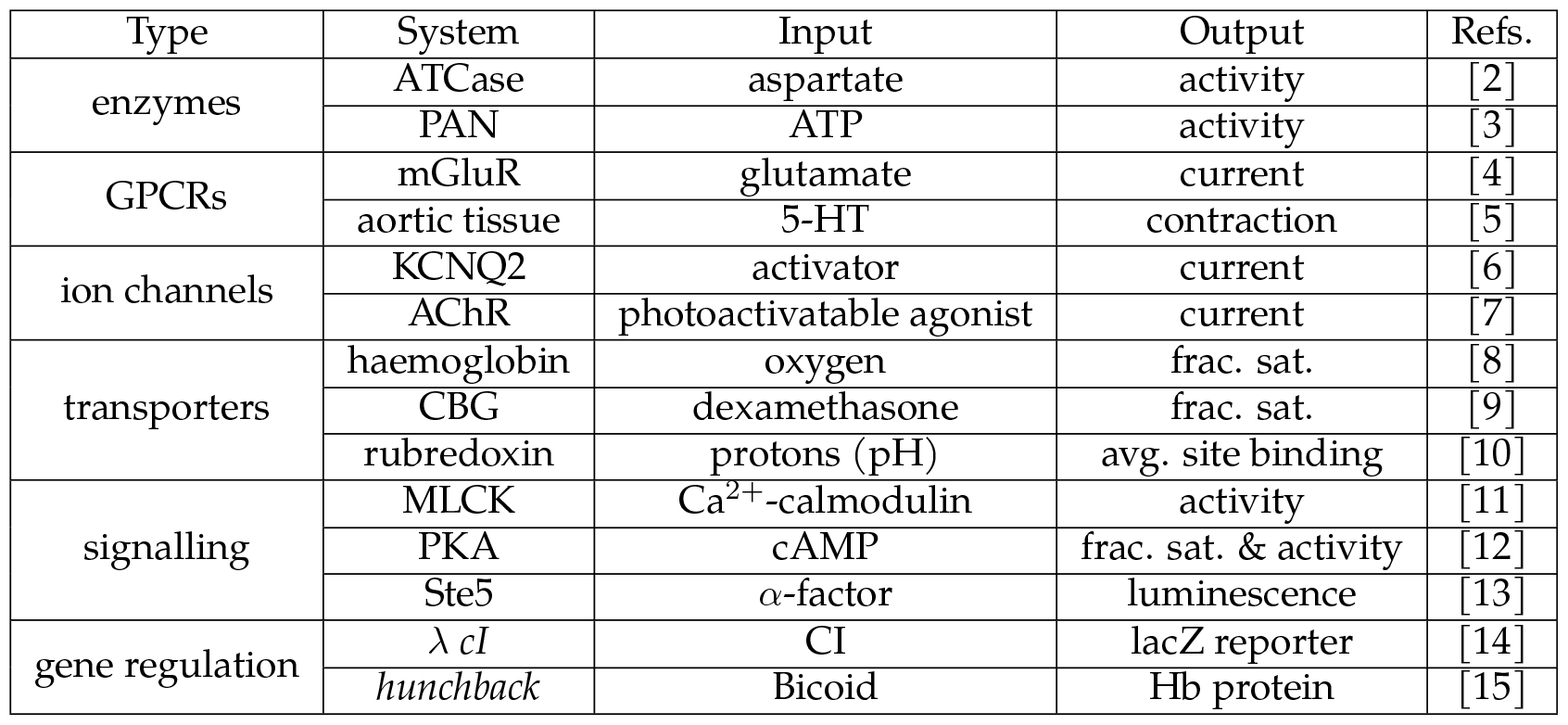
Experimentally measured input-output responses. For the meaning of an acronym for a system, see the corresponding citation. Avg.: average; frac. sat.: fractional saturation.

## 2 Examples of linear framework graphs

Linear framework graphs can represent different forms of molecular complexity [16], as illustrated in Fig.S1. The product construction in Fig.S1A shows how the hypercube structure, *C*_*m*_, can be defined as

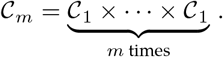

The product construction can be applied to any two graph structures; see [16] for details. Hierarchical nesting, which arose in studying allostery [18], is another procedure, akin to the product construction, that can generate increasingly complex graph structures.

## 3 Eq. 9 gives all input-output responses at t.e

Suppose given a rational function *r*(*x*) of degree *l*, defined as in Eq. 9 of the main text with the accompanying restrictions on the coefficients *α*_*i*_ and *β*_*i*_. We show here that labels can be assigned to the graph structure 𝒞_*l*_ so that it can reach t.e., with *x* being the concentration of the unique binding ligand, and coefficients *λ*_*i*_ can be found, as in Eq.5 of the main text, such that *r*(*x*) is the resulting input-output response.

**Figure S1:**
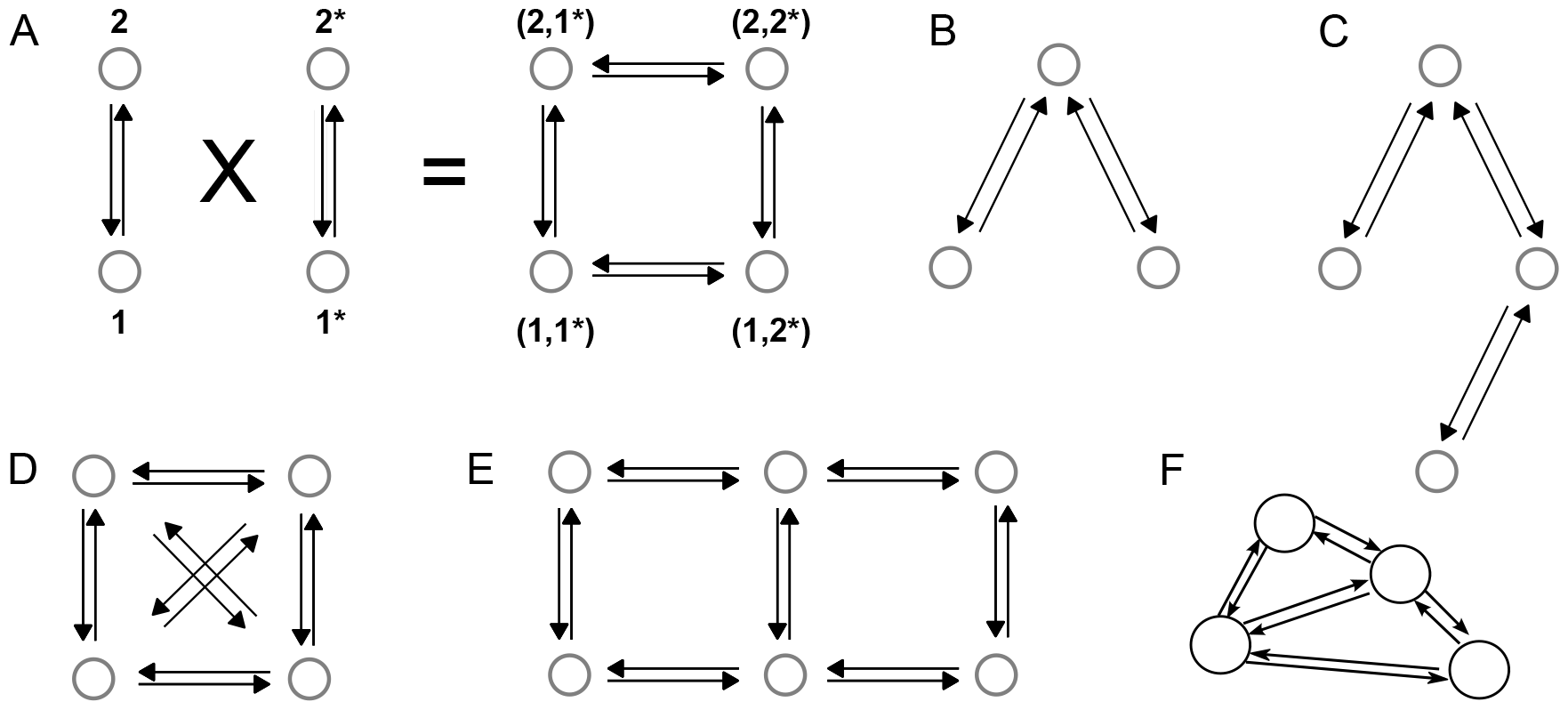
Graph structures; labels are omitted for clarity. **A**. Product construction, with the vertices indexed to illustrate how the hypercube structure, 𝒞_2_, arises as the product of 2 copies of 𝒞_1_. **B**. Graph structure that could represent mutual exclusion, such as two binding sites which are sufficiently close together that only one site can be bound at a time. **C**. Graph structure that could represent ordering, such as two binding sites in which binding first to site 1 allows binding to site 2 but binding first to site 2 excludes binding to site 1. **D**. Graph structure that could represent 4 options which can arbitrarily interchange, such as methylation patterns on a lysine residue. **E**. Lattice graph structure in which the vertices could represent conformations or discrete spatial locations. **F**. Graph structure representing the conformational ensemble of the pentameric, ligand-gated ion channel GLIC, constructed by Markov State Modelling, adapted from [17].

There is a standard labelling for hypercube structures at t.e. that was introduced in [19, SI, §3], which should be consulted for more details. Set theory notation is needed to explain this labelling. Each vertex of the hypercube structure corresponds to a subset of bound sites. Let *S* ⊆ {1, *· · ·, l*} denote that subset and use this symbol as the index for that vertex. The edges of *C*_*l*_ are then given by *S* ⇋ *S* ∪ {*i*} where *I* ∉ *S*. At t.e., only the label ratios are needed to determine s.s. probabilities. Let the association constant, *K*_*i,S*_, denote the ratio,

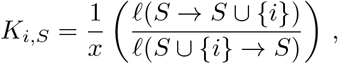

so that *xK*_*i,S*_ is the label ratio. The *K*_*i,S*_ have dimensions of (concentration)^−1^ and there are *l*2^*l*−1^ such association constants. Whatever unit is used for concentration, we will assume it is the same one used in Eq.9 in the main text. Because of the cycle condition, the association constants are not independent.

Restricting to those *K*_*i,S*_ for which *i* is less than all the sites in *S*, denoted *i < S*, yields an independent and complete set of 2^*l*^ − 1 parameters for *C*_*l*_ at t.e. Note that, since the empty set, ∅, has no members, the condition *i <* ∅ is trivially satisfied for all sites *i* and so the association constant, *K*_*i*,∅_, for binding to site *i* with no other sites bound, is one of the independent parameters. For convenience, we will refer to the resulting graph as 𝒞_*l*_.

In terms of this parameterisation, the vector *µ*(𝒞_*l*_) is given as follows. Suppose that *S* = {*i*_1_, *· · ·, i*_*k*_} is a vertex of 𝒞_*l*_ for which the sites are written in order, so that *i*_1_ *< i*_2_ *< · · · < i*_*k*_. Then, the vertex *S* may be reached from the empty subset, ∅, by a path of reversible edges such that,

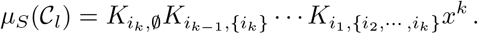

By construction, these association constants are included in the independent set of parameters. It is now straightforward to determine the s.s. probability of vertex *S* from Eq.4 of the main text. It follows that the denominator in Eq.4 of the main text is given by

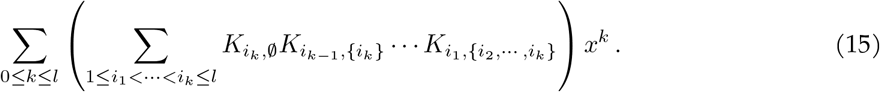

The term in Eq.15 with *k* = 0, for which there is an empty product between the brackets, is taken by convention to be 1.

Returning to the rational function *r*(*x*) in Eq.9 of the main text, we may assume, by dividing above and below by *β*_0_, that *β*_0_ = 1. There are *l* remaining coefficients in the denominator, *β*_1_, *· · ·, β*_*l*_, which are given. Since there are 2^*l*^ − 1 association constants, which exceeds *l* as soon as *l >* 1, it is straightforward to recursively choose them to yield the correct denominator polynomial.

We will illustrate this for *l* = 3, from which the reader should be able to readily see the general procedure. We need to give values to the 7 = 2^3^−1 independent association constants, *K*_1,∅_, *K*_2,∅_, *K*_3,∅_, *K*_1,{2}_, *K*_1,{3}_, *K*_2,{3}_ and *K*_1,{2,3}_, and choose the 8 = 2^3^ *λ*_*S*_ coefficients for the input-output response. Comparing Eq.15 to the denominator of Eq.9 in the main text, we first require that

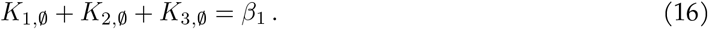

Choose *K*_1,∅_ and *K*_2,∅_ arbitrarily so that 0 *< K*_1,∅_ + *K*_2,∅_ *< β*_1_, which we may always do, and then choose *K*_3,∅_ to satisfy Eq.16. Next we require that

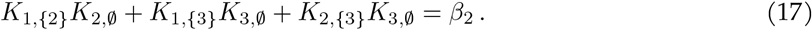

As before, we may choose *K*_1,{2}_ and *K*_1,{3}_ arbitrarily so that,

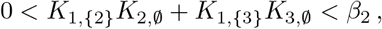

which we may again always do. The key point here, which makes the recursion work, is that the quantities to be chosen satisfy a linear equation (Eq.17) in which the previously chosen quantities give the coefficients. We may now choose *K*_2,{3}_ to satisfy Eq.17. Finally, we require that

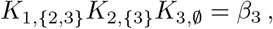

which we may easily satisfy by choosing *K*_1,{2,3}_ = *β*_3_/(*K*_2,{3}_*K*_3,∅_). This determines the 7 association constants. Now choose *λ*_∅_ = α_0_, *λ*_{1}_ = *λ*_{2}_ = *λ*_{3}_ = α_1_*/β*_1_, *λ*_{1,2}_ = *λ*_{1,3}_ = *λ*_{2,3}_ = α_2_*/β*_2_ and *λ*_{1,2,3}_ = α_3_*/β*_3_. Because of the restriction that 0 ≤ *α*_*i*_ ≤ *β*_*i*_, as stipulated for Eq.9, it follows that 0 ≤ *λ*_*S*_ ≤ 1, as required for Eq.5. It is not difficult to see that we recover *r*(*x*) from these choices and that a similar procedure works in general for any *l >* 1. This completes the proof.

## 4 Intrinsic measures of sharpness

The rational functions defined b y Eq.9 c an b e s ubstantially m ore c omplicated t han s igmoidal inputoutput responses. They may have positive basal values at *x* = 0 and be non-monotonic with multiple peaks and troughs and they may asymptote as *x* → ∞ to a value other than their maximum (Fig.S2). Non-monotonic responses occur in areas like pharmacology and toxicology; particularly intricate examples are found in pH titrations of biomolecules [10]. Fig.S2 shows an example based on Eq.9 of the main text, which also illustrates how the normalisation value, *x*_0.5_, in Eq.10 of the main text is chosen.

**Figure S2:**
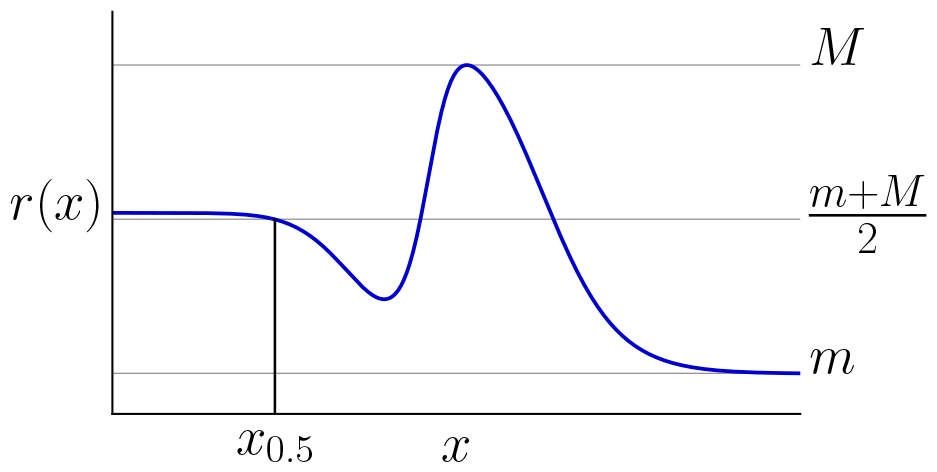
Rational function and normalisation. The plot shows an example input-output response, *r*(*x*), obtained from Eq.9 for *l* = 6, with the following coefficients, α_0_ = 5, *α*_1_ = 50, *α*_2_ = 0.1, *α*_3_ = 10, *α*_4_ = 0.01, *α*_5_ = 1, *α*_6_ = 0.001, *β*_0_ = 10, *β*_1_ = 100, *β*_2_ = 100, *β*_3_ = 10, *β*_4_ = 0.01, *β*_5_ = 1, *β*_6_ = 0.01.

We asserted in the main text that the supremum of |*dq/dy*| is attained for a finite value of *y*. To see this, note from the structure of the rational function *r*(*x*) in Eq.9 of the main text that *dq/dy* = *dr/dx · x*_0.5_ must asymptote to the *y*-axis at infinity, lim_*y*→∞_ *dq/dy* = 0. Hence, if the supremum of |*dq/dy*| is only found asymptotically when *y* → ∞, it must be the case that *dq/dy* = 0 for all *y* ∈ [0, ∞). Hence, *q*(*y*) is a constant and therefore so too is *r*(*x*). But this is impossible because the input ligand must bind to at least one site, so that the degree *l* in Eq.9 satisfies *l* ≥ 1 and *r*(*x*) cannot be a constant.

## 5 Estimating position and steepness

Numerical estimation of position and steepness was carried out differently depending on whether or not the underlying model is at t.e. For an equilibrium model, the coefficients of the rational function in Eq.9 of the main text can be algebraically calculated. Away from equilibrium, this is no longer feasible because of the combinatorial explosion described in the main text. Instead, s.s. probabilities were estimated directly from the Laplacian matrix in Eq.13 of the main text by singular value decomposition. The code for (*p, s*) calculations is available in the GitHub repository github.com/rosamc/universal-boundaries-Hopfi eld-barrier.git. The ranges and densities of points used here were determined by trial-and-error, to the point where further increases did not significantly change (*p, s*) values.

### 5.1 Equilibrium models

To calculate (*p, s*) values for equilibrium models quickly and accurately, we implemented a custom algorithm in C++, called from Python using `pybind11`, using high-precision floating-point types provided by the GNU MPFR library through the Boost interface (www.boost.org). After extensive testing, we found similar results for 50 or 100 digit precision, so 50 digit precision was used. The algorithm works as follows.

Depending on how a particular model is parameterised, which is discussed further below, the coefficients of the rational function *r*(*x*) in Eq.9 are calculated in terms of these parameters. Unless (*m*(*r*)+*M* (*r*))/2 = 0.5, which is the case for the fractional saturation input-output response for Model I (below), *r*(*x*) is evaluated at the points *x* = 10^−60+*k*.0.02^ for *k* = 0, 1, *· · ·*, 5999. The quantities *m*(*r*) and *M* (*r*) in Eq.10 are estimated and *x*_0.5_ is determined by finding the smallest positive solution to the polynomial equation, *r*(*x*) = (*m*(*r*) + *M* (*r*))/2. Polynomial solving was performed with a custom C++ implementation of the Aberth-Ehrlich root-finding method [20, 21]. The code returns complex roots. Roots having a positive real part, and an imaginary part with absolute value smaller than 10^−15^ are considered to be real zeros, while the other roots are discarded. This polynomial solving code is available at github.com/kmnam/p olynomials.git.

Having calculated *x*_0.5_, the normalised function *q*(*y*) = *r*(*yx*_0.5_) is obtained by multiplying *α*_*i*_ and *β*_*i*_ in Eq.9 by (*x*_0.5_)^*i*^. The derivatives *dq/dy, d*^2^*q/dy*^2^ and *d*^3^*q/dy*^3^ are then calculated algebraically. The roots of *d*^2^*q/dy*^2^ are estimated by polynomial solving, using the algorithm just described. The local maxima and minima of *dq/dy* are determined by evaluating the sign of *d*^3^*q/dy*^3^ at each root and the maximum of |*dq/dy*| for *y* ∈ [0, ∞) is determined. Eq.11 of the main text then gives the (*p, s*) values.

### 5.2 Non-equilibrium model in Fig. 3C

Because of the combinatorial explosion away from t.e., as discussed in the main text, it is not computationally feasible to determine the coefficients of the rational function *r*(*x*) in Eq. 9 in terms of the model parameters. We therefore numerically evaluated the input-output response and estimated (*p, s*) values using finite differences.

**Figure 3:**
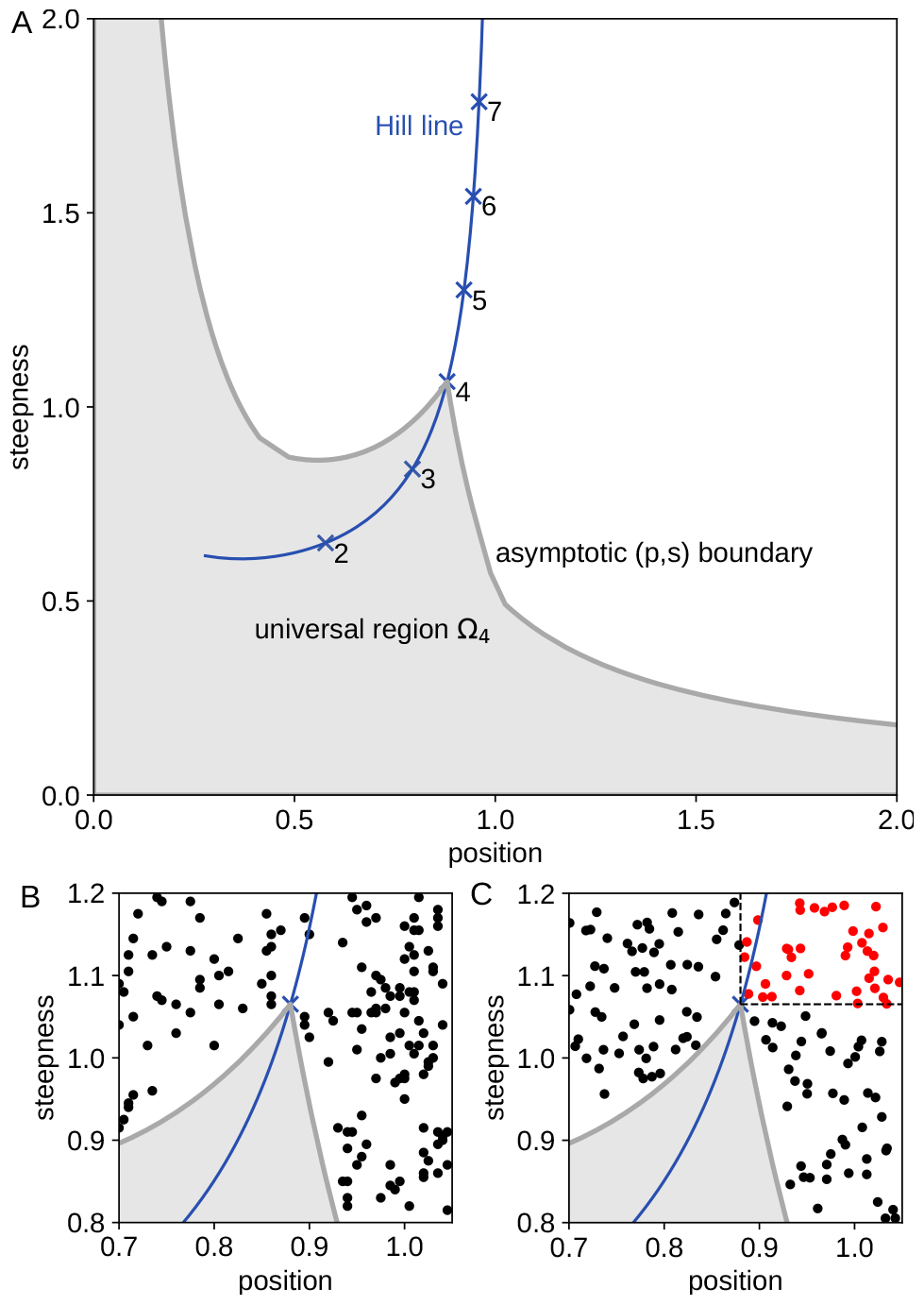
Universal position-steepness region and the Hopfield barrier. **A**. Universal asymptotic (*p, s*) region, Ω_4_ (gray area), for thermodynamic equilibrium models with *m* = 4 binding sites for the input ligand, with the Hill line shown as in Fig.1C. **B**. Expanded view of the cusp showing (*p, s*) points lying outside the universal region (black dots) for rational functions which do not satisfy the coefficient condition, α_*i*_ ≤ *β*_*i*_, for Eq.9. Only points outside Ω_4_ are shown for clarity. **C**. Expanded view of the cusp showing (*p, s*) points lying outside the universal region for the graph with the hypercube structure *C*_4_, output given by fractional saturation and parameter values chosen away from thermodynamic equilibrium. The red points beyond the dashed lines exceed ℋ_4_ in both position and steepness and confirm that the Hill function *ℋ*_4_ is the Hopfield barrier for the sharpness of models with *m* = 4 input binding sites. Only points outside Ω_4_ are shown for clarity.

To calculate *r*(*x*) at a given *x* we used singular value decomposition (SVD) directly on the matrix ℒ (*G*), described in Eq.13 of the main text. SVD gives a basis for ker ℒ (*G*), from which the s.s. probabilities can be calculated, as in Eq.4 of the main text, and *r*(*x*) thereby determined. This was again done in C++ using 100-digit precision floating-point types provided by the MPFR library through the Boost interface, with the BDCSVD routine from the Eigen library. This code is available in github.com/rosamc/universal-b oundaries-Hopfield-barrier.git.

For accuracy, it is necessary to evaluate *r*(*x*) over a large range and at many values of *x*, which makes the calculation slow. To balance computation time against accuracy, we followed a two-step approach. First, we get an initial estimate of *x*_0.5_. The rational function *r*(*x*) is sparsely evaluated at the points *x* = 10^−60+*k×*1.62^ for *k* = 0, *· · ·*, 74. Then we use the `interp1d` function in the Python Scipy library to interpolate values of *r*(*x*) at *x* = 10^−60+*k×*0.0012^ for *k* = 0, *· · ·*, 99999. We use these interpolated values to estimate *m*(*r*), *M* (*r*) and *x*_0.5_ using Eq. 10 in the main text.

From this initial estimate of *x*_0.5_, we re-evaluate *r*(*x*) over a smaller range more densely. We calculate *r*(*x*) at 10^3^ logarithmically-spaced points between 10^−4^ *× x*_0.5_ and 10^3^ *× x*_0.5_ and interpolate values of *r*(*x*) at 10 logarithmically-spaced points within this range, 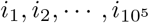. We then calculate a more accurate *x*_0.5_. The value of |*dr/dx*| is calculated by finite differences as |(*r*(*i*_*k*+1_) − *r*(*i*_*k*_))/(*i*_*k*+1_ − *i*_*k*_)| for *k* = 1, *· · ·*, 10^5^ − 1 and the maximum value and corresponding input point, *i*_*m*_, are obtained. This gives the un-normalised steepness *s*^*u*^(*r*), and the un-normalised position, *p*^*u*^(*r*), as defined in the main text, from which the (*p, s*) values are calculated by *p*(*r*) = *p*^*u*^(*r*)*/x*_0.5_ and *s*(*r*) = *s*^*u*^(*r*)*x*_0.5_.

## 6 Numerically estimating (*p, s*) regions

We explain here in more detail the procedure used to estimate the universal (*p, s*) regions Ω_4_ (Fig.3A and Fig.S4) and Ω_6_ (Fig.S5) and the (*p, s*) regions for various specific models (Fig.1C, Fig.S6). The basic idea is to use iterative, biased sampling of the relevant parameters, whose values are drawn from a parametric range [10^−*a*^, 10^*a*^], to reach a boundary for that value of *a*. Boundaries are calculated for increasing values of *a* and an asymptotic boundary is considered to have been reached when the boundaries for two consecutive *a* values coincide when plotted together; see Fig.S4 for Ω_4_ and Fig.S5 for Ω_6_. Experience has shown that the algorithm can get trapped prematurely and the procedures outlined below have been refined by trial-and-error specifically to overcome this problem. The method was originally developed in [19] and subsequently elaborated in [22]. The Python code for the boundary procedure is available in the GitHub repository github.com/rosamc/GeneRegulatoryFunctions.git.

### 6.1 Generalities

(*p, s*) regions are estimated within some rectangular box within the positive quadrant, which determines the extent of (*p, s*) space to be explored. Unless otherwise noted, position was explored from 0.4 to 2.5, and steepness from 0.3 to 2.5. The box is divided into a grid of square “cells” of width and height 0.005 in position and steepness. (*p, s*) regions are extended in an iterative manner from an initial seed region. A grid cell is considered to be filled when a (*p, s*) point falls within it; a filled cell is on the boundary of the current region if one of its four immediate neighbours (above, below, left or right) is not filled.

The parametric range for the relevant parameters is chosen as [10^−*a*^, 10^*a*^] for some value of *a >* 0. The sequence of *a* values used to establish an asymptotic boundary depends on the (*p, s*) region being estimated and is given in Table S2. The relevant parameters may involve constraints, which are always in the form of inequalities. For example, for the universal region Ω_*m*_, the relevant parameters are the coefficients, *α*_0_, *· · ·, α*_*m*_ and *β*_0_, *· · ·, β*_*m*_, of the rational function in Eq.9, and these are subject to the constraint that *α*_*i*_ ≤ *β*_*i*_. Other parametric constraints arise for some of the models described below. We will refer to those parameters, like *β*_*i*_, that lie within a semi-infinite interval, as “independent”, and the others, like *α*_*i*_, whose range is limited to a finite interval by the independent parameters, as “dependent”. The search starts by randomly sampling parameter sets from the specified range, ensuring the constraints are satisfied, until 10 grid cells are filled (see Initialisation below). Only one parameter set is kept for each filled cell. The boundary is subsequently extended by modifying the parameter sets within boundary cells and iteratively exploring the parameter space to grow the region until the final boundary is reached. The algorithm described below does not necessarily give rise to a boundary that is always a simple closed curve, with a clearly defined inside and outside, but we encountered no difficulties in this respect in practice.

During the boundary estimation, only the grid cell corresponding to a (*p, s*) point is recorded, alongside the corresponding parameter set, not the exact (*p, s*) value. Once the final boundary is reached, Mathematica is used to recalculate the (*p, s*) values for those parameter sets on the boundary, thereby providing an independent test of the final boundary. The Mathematica algorithm paralleled the one described previously: definition of *r*(*x*) from the parameter values, evaluation over [10^−60^, 10^60^], calculation of *x*_0.5_, definition of *q*(*y*), calculation of derivatives, calculation of the extrema of *dq/dy*. The Mathematica function `Solve` was used to determine the roots of polynomials and the function `D` was used to calculate derivatives. The Mathematica code is available in github.com/rosamc/universal-boundaries-Hopfi eld-barrier.git.

A (*p, s*) point calculated by Mathematica is retained to calculate the definitive boundary only when it falls in the original grid cell for the corresponding parameter values. Otherwise, the point is discarded. In the vast majority of cases there was excellent agreement, with most of the discrepancies arising when the Mathematica-calculated point was in a grid cell next to the original one (see Further details below).

The search process is delimited by hyperparameters. Some of them are fixed throughout: *niters_conv* = 1500; *niters_conv_points* = 1000; *niters_target* = 500; *L_project* = 15; *tol_target* = 0.001. Some of them are varied between runs: *prob_par* = 0.2, 0.5; *prob_replace* = 0.2, 0.6; *extremesu* = [−2, 2], [−1.5, 1.5], [−1, 1], [−0.5, 0.5]. (The meaning of these hyperparameters is given in the description of the search algorithm below.) We run the boundary estimation for each of the 16 = 2 *×* 2 *×* 4 combinations of the three varying hyperparameters, starting in each case from a different seed region (below) to avoid trapping. There were differences in the rate of convergence, with a few, rare cases failing to converge in the wall time allowed, or experiencing cluster-related impediments (Table S2). For those that converged, we never noticed any substantial difference among the boundaries computed with different hyperparameter combinations, suggesting the boundary has been accurately estimated in each case. In order to estimate the final boundary, we took the parameter sets corresponding to boundary cells for each of the converged runs and calculated their (*p, s*) value using Mathematica as explained above. The points where the Mathematica-calculated (*p, s*) point fell in the original grid cell were collected together and the definitive boundary determined from this collection obtained from all converged runs (in which some cells may no longer be on the boundary). To draw the definitive boundary shown in the figures, the `alphashape` routine of the Python Alpha Shape toolbox (https://pypi.org/project/alphashape) was used, with a boundary-specific, manually-adjusted alpha parameter to ensure a good match to the input (*p, s*) points. The C++ code, Mathematica code and the Python Jupyter notebooks used to determine the boundaries and make the figure plots are available in github.com/rosamc/universal-boundaries-Hopfield-barrier.git.

### 6.2 Initialisation

Having defined the grid and specified the search hyperparameters, as outlined above, the first step is to identify a few parameter sets whose (*p, s*) values fall within different grid cells. For this, parameter values are randomly chosen from the specified parametric range, subject to the appropriate inequality constraints. For example, for the universal models, the log_10_ *β*_*i*_ are chosen uniformly at random within the range [−*a, a*], and the log_10_ α_*i*_ are then chosen uniformly at random in the range [−*a*, log_10_ *β*_*i*_], so as to satisfy the constraint α_*i*_ ≤ *β*_*i*_. Constraints for other models are treated similarly. A (*p, s*) value is calculated as described previously and the corresponding grid cell is considered filled. If a (*p, s*) value falls into an already filled cell, the new parameter set is discarded and the old one retained. The process is repeated until 10 different cells are filled and the corresponding current working boundary is defined as above.

### 6.3 Extension

After initialisation is completed, the current working boundary is iteratively extended until it stabilises. The extension procedure involves slightly changing the parameter sets of the cells on the current working boundary, with the intention that these new parameter sets will yield slightly different (*p, s*) values, hopefully corresponding to empty cells outside the current working boundary. At each iteration, the parameter sets are changed in different ways in order to explore their neighbourhood, through the *Mutation* and *Pulling* steps explained below. The boundary is assumed to have stabilised when no new boundary cell is filled for *niters_conv* consecutive iterations. If a cell remains at the boundary for *niters_conv_points* consecutive iterations, then it is considered to be on the final boundary and the *Mutation* and *Pulling* steps are omitted for that cell. This reduces the computational cost of the procedure in case the boundary converges fast in a given region but not another.

#### Mutation

At each iteration, new parameter sets are generated and tested for their ability to produce a (*p, s*) point that fills a cell that is either outside or at the current boundary. If the filled cell is at the current boundary, the old parameter set is replaced by the new one with probability *prob_replace*. This stochastic replacement of boundary points helps to escape local trapping: by slightly changing parameter sets on the boundary, it may become possible to move outside the boundary in subsequent iterations. For each boundary parameter set, the algorithm attempts at most 20 trials to modify its value and generate a new accepted parameter set, whose cell is either outside or at the current boundary. Trials are halted as soon as a new parameter set is accepted. After applying the mutation procedure to all boundary parameter sets, the current working boundary is recomputed.

To generate a new parameter set from an old one, each parameter value *v* is replaced by *z* with probability *prob_par*, where *z* is obtained from *v* in a different way depending on the trial number, as specified below. The hyperparameter *prob_par* stochastically controls by how much a given parameter set is changed: the larger value of 0.5 is suitable for quick exploration, while the smaller value of 0.2 helps to fine tune the boundary near stabilisation. Let the value of the hyperparameter *extremesu* be [fmin, fmax] (with fmin *<* 0, fmax *>* 0), and let 𝒩 (*µ, σ*) be the normal distribution with mean *µ* and standard deviation *σ*.

- trials 1-2: choose *θ* randomly from the uniform distribution on [fmin, fmax] and set *z* = *v ×* 10^*θ*^.
- trials 3-5: choose *z* randomly from the normal distribution 𝒩 (*v, v/*0.1);
- trials 6-9: choose *z* randomly from the normal distribution 𝒩 (*v, v/*0.5);
- trials 10-13: choose *z* randomly from the normal distribution 𝒩 (*v, v*);
- trials 14-15: choose *z* randomly from the normal distribution 𝒩 (*v, v/*2);
- trials 16-20: choose *θ* randomly from the uniform distribution on [fmin/0.5, fmax/0.5] and set *z* = *v ×* 10^*θ*^.

**Table S2:** Details of the boundary estimation runs. The columns record the type of model (Model); the parametric range exponent, *a*, in [10^−*a*^, 10^*a*^] (Range); the number of successfully completed runs (# Runs); the number of those completed runs in which all boundary points calculated by Mathematica were in the original cell (# Correct); the number of completed runs with less than 2% of boundary points in a neighbouring cell to the original ((0 − 2)%); the number of completed runs with at least 2% and no more than 5% of boundary points in the neighbouring cell ([2 − 5)%); some other possibility that is explained in the Notes to this caption (Other). For the various parts of Ω_4_, see Fig.S3. The *a* = 10 analysis for Model IV was exceptionally slow and failed to converge. As there was little difference between the regions for *a* = 7 and *a* = 9, *a* = 10 was abandoned. Notes. (†) — there were 13 runs with [0.2 − 2.2)% of points for which Mathematica was unable to find the roots of the polynomials; 3 runs with [0.2 − 0.5)% of non-neighbouring cell points; 13 runs with [9 − 15)% neighbouring cells; and 3 runs with [15 − 18)% neighbouring cells. (‡) — there were 12 runs with [0.1−1)% of points for which Mathematica was unable to find the roots of the polynomials; and 5 runs with [0.1 − 0.2)% of non-neighbouring cell points.

| Model | Range | # Runs | # Cor-rect | # (0-2)% | # [2-5)% | Other |
| --- | --- | --- | --- | --- | --- | --- |
| $\Omega_4$<br>(Fig.3) | 0.3 | 16 | 9 | 7 | 0 | - |
|  | 0.5 | 16 | 0 | 16 | 0 | - |
|  | 0.7 | 16 | 0 | 16 | 0 | - |
|  | 1 | 16 | 0 | 16 | 0 | - |
|  | 2 | 16 | 0 | 15 | 1 | - |
|  | 3 | 16 | 0 | 15 | 1 | - |
| lower right | 3 | 16 | 0 | 0 | 0 | † |
| upper left | 3 | 16 | 9 | 7 | 0 | - |
| lower left | 3 | 16 | 0 | 16 | 0 | ‡ |
| $\Omega_6$ (Fig.S5) | 0.3 | 16 | 0 | 16 | 0 | - |
|  | 0.5 | 16 | 0 | 3 | 13 | - |
|  | 0.7 | 12 | 0 | 1 | 11 | - |
|  | 1 | 11 | 0 | 1 | 10 | - |
|  | 2 | 16 | 0 | 6 | 10 | - |
|  | 3 | 16 | 0 | 2 | 14 | - |
| I (Fig.S6) | 1 | 16 | 16 | 0 | 0 | - |
|  | 2 | 16 | 16 | 0 | 0 | - |
|  | 3 | 16 | 16 | 0 | 0 | - |
|  | 5 | 15 | 15 | 0 | 0 | - |
|  | 7 | 16 | 16 | 0 | 0 | - |
| II (Fig.S6) | 0.5 | 14 | 14 | 0 | 0 | - |
|  | 1 | 15 | 15 | 0 | 0 | - |
|  | 2 | 16 | 16 | 0 | 0 | - |
|  | 3 | 16 | 16 | 0 | 0 | - |
|  | 6 | 16 | 16 | 0 | 0 | - |
| III (Fig.S6) | 1 | 16 | 0 | 4 | 12 | - |
|  | 2 | 16 | 0 | 9 | 7 | - |
|  | 3 | 16 | 0 | 11 | 5 | - |
|  | 4 | 16 | 0 | 12 | 4 | - |
| IV (Fig.S6) | 3 | 16 | 16 | 0 | 0 | - |
|  | 5 | 16 | 16 | 0 | 0 | - |
|  | 7 | 16 | 16 | 0 | 0 | - |
|  | 9 | 16 | 16 | 0 | 0 | - |
|  | 10 | 0 | N/A | N/A | N/A | - |
| V (Fig.S6) | 0.5 | 16 | 16 | 0 | 0 | - |
|  | 1 | 16 | 16 | 0 | 0 | - |
|  | 2 | 16 | 16 | 0 | 0 | - |
|  | 3 | 16 | 15 | 1 | 0 | - |
|  | 6 | 16 | 12 | 4 | 0 | - |
| VI (Fig.S6) | 0.5 | 16 | 16 | 0 | 0 | - |
|  | 1 | 15 | 15 | 0 | 0 | - |
|  | 2 | 16 | 16 | 0 | 0 | - |
|  | 3 | 15 | 15 | 0 | 0 | - |
|  | 6 | 16 | 16 | 0 | 0 | - |

Note that this procedure can result in parameter values outside the allowed parametric range [10^−*a*^, 10^*a*^] or the range determined by the constraints, as described above. If a parameter choice *z* lies outside the desired parameter range [*c, d*], then the nearest value to *z* on the boundary of the range is chosen instead: if *z < c < d*, then *c* is chosen, while if *c < d < z*, then *d* is chosen. This can be important to find parameter combinations that lead to extreme regions of the (*p, s*) space. If there are constraints among the parameters, then the independent parameters, as defined above, are chosen first and the dependent parameters chosen afterwards. If a dependent parameter is not selected for change because of the stochasticity associated with *prob_par* but one of the independent parameters that determines its constraint is selected, the dependent parameter may no longer satisfy its constraint. If so, this dependent parameter is changed by the smallest amount that allows its constraint to be satisfied.

#### Pulling

After applying the *Mutation* step, a different strategy is used of “pulling” towards a target grid cell, *t*, that is chosen for each boundary cell *c*, so as to lie outside the current boundary. Each cell is assigned integer coordinates based on its position in the grid. If the grid has *K* rows and *L* columns, the coordinates are the corresponding row and column numbers chosen in [0, *K* − 1] *×* [0, *L* − 1]. Two kinds of targets are defined, one to pull away from the boundary and one to pull away from the centre of the region.

For the first target, the nearest boundary cell, *b*, is identified that is no more than 3 cells to the right of *c*. This target is only used if there are more than 100 cells at the boundary in order to have a good chance of finding *b*. If no such cell exists, *c* is skipped. If *b* is found, the line joining the integer coordinates of *c* and *b* is rotated around the coordinates of *c* by an angle of *π/*4 away from the current boundary. The cell that lies farthest on this line, while remaining within the grid and no farther than *L_project* cells away, is chosen as *t*.

For the second target, the centroid, ζ, of the filled cells is calculated. The line joining the integer coordinates of ζ and *c* is extended away from the boundary. The cell that lies farthest on this line from *c*, while being outside the boundary and remaining within the grid and no farther than Euclidean distance *L_project*, is chosen as *t*.

Having chosen a target cell *t* for a given boundary cell *c*, the exploration proceeds as follows. A new parameter set is generated from the parameter set for *c*, as described above for trial 1 of the mutation step. If the Euclidean distance from the (*p, s*) of this new parameter set to the bottom left corner (*p, s*) of *t* is reduced, the old parameter set is replaced with the new one. If not, the old parameter set is retained and used again. The process is iterated until either the new distance to *t* is less than *tol_target* or the number of iterations reaches *niters_target*. If the resulting parameter set fills a new cell, the boundary will have been extended. After visiting all cells on the current boundary, a new working boundary is computed.

### 6.4 Further details

Boundary estimation for the universal Ω_4_ region in Fig. 3 was done in parts to reduce the memory requirements (Fig. S3). We first identified the parametric range corresponding to the asymptotic boundary for the central region, we then extended the “wings”, and we finally explored the lower-left region of the quadrant (Table S2). Let [*c, d, f, g*] denote the rectangular box between position values [*c, d*], where *c < d*, and steepness values [*f, g*], where *f < g*. The boundary lying in the central box [0.4, 2.5, 0.3, 2.5] was estimated with cells of size 0.005, as described above. The boundaries for the parametric ranges [10^−2^, 10^2^] and [10^−3^, 10^3^] overlapped, so we took the latter range as the one corresponding to the asymptotic boundary. Then, we estimated the “wings” lying in the boxes [1.2, 2.1, 0, 0.4] (lower right) and [0, 0.5, 0.8, 2.1] (upper left), using the same range of [10^−3^, 10^3^]. For each of these two wing boxes, we seeded the search with the boundary points identified in one of the runs for the central box, which fell within the respective wings. Finally, to double check this extension and ensure that we could extend the boundary down to small (*p, s*) values, we also estimated the boundary in the box [0, 1.5, 0, 1.8] (lower left), using a coarser grid of cells of size 0.01, again to reduce the memory cost of the calculation.

**Figure S3:**
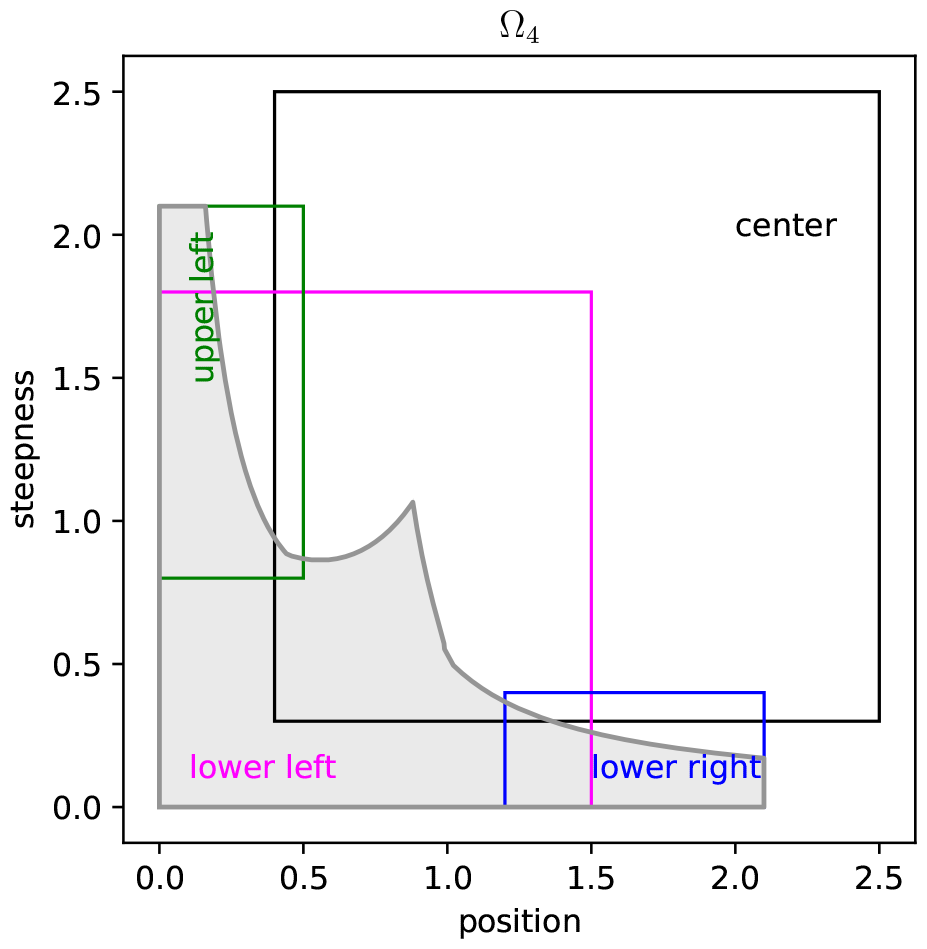
Grid boxes used to estimate the universal region Ω_4_, as described in the text.

Table S2 gives details of the boundary estimation for the various models considered in the paper. As noted above, cluster-related issues sometimes led to unfinished runs or the wall time allocated was insufficient. Since the majority of the jobs finished, we did not repeat the searches for the few that did not. Similarly, in some cases, the (*p, s*) values calculated by Mathematica for boundary parameter sets did not fall in the same cell (Table S2). These parameter sets were discarded and not used to calculate the final boundary. It can be seen from Table S2 that when Mathematica values are non-coincident, the corresponding (*p, s*) values nearly always fall in a neighbouring cell, with the exceptions being only for the lower-right and lower-left boxes of Ω_4_. For the universal models, Ω_4_ and Ω_6_, the non-coincident points are always far from the cusp on the Hill line, typically in regions of high position and low steepness. For Model III, in contrast, the non-coincident points are on the boundary of the region, at high values of steepness, but these points are always far from the boundary of Ω_4_.

**Figure S4:**
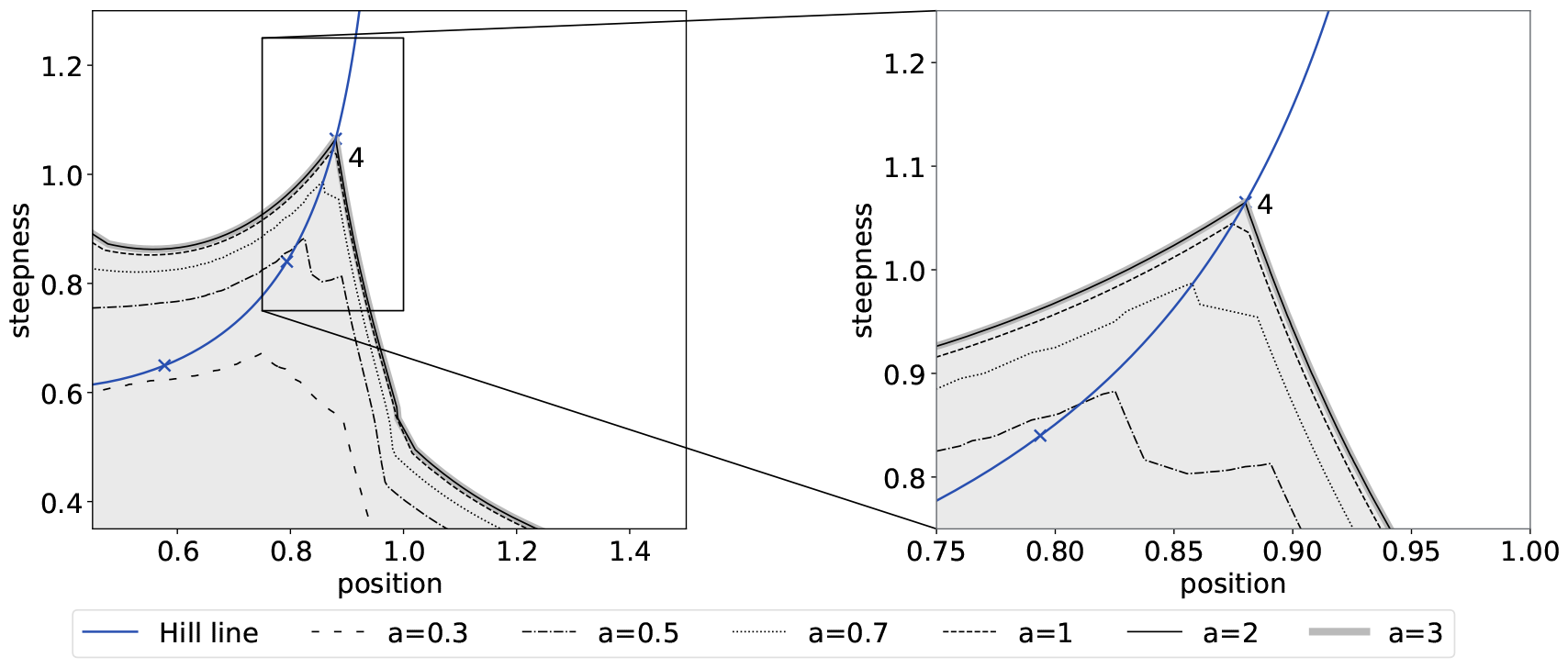
Universal region Ω_4_, showing convergence to the asymptotic boundary for increasing parametric ranges, [10^−*a*^, 10^*a*^], shown as varying linestyles, as indicated in the key. The same conventions are followed as in Fig.3A of the main text, with the (*p, s*) value for ℋ_4_ marked. The right-hand panel shows in greater resolution the box around the cusp in the left-hand panel.

**Figure S5:**
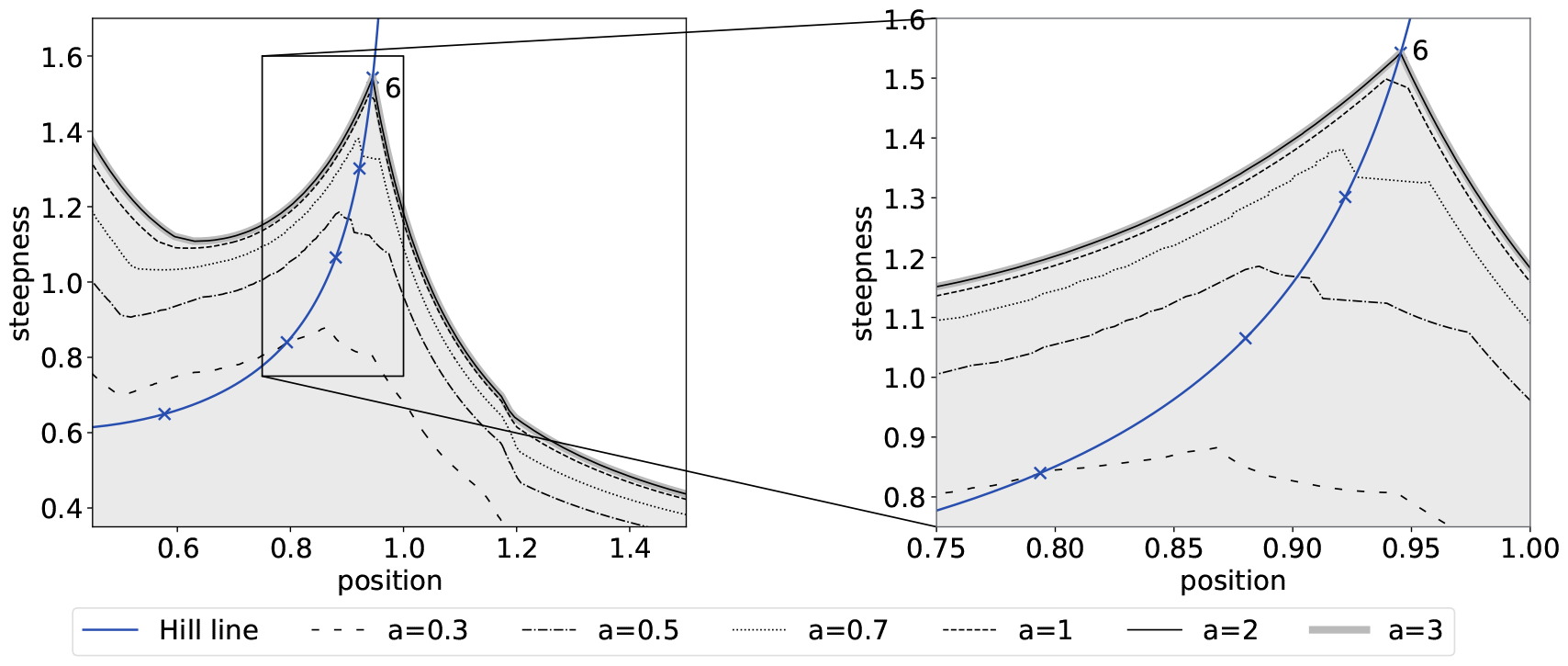
Universal region Ω_6_, showing convergence to the asymptotic boundary, following the same format as in Fig.S4.

## 7 Six specific models

As a partial test of the universal (*p, s*) region Ω_4_, we estimated the asymptotic region for six previously-studied models at t.e., each with 4 input binding sites, and confirmed that they all fell within Ω_4_ (Fig.S6). In the descriptions which follow, we use *G* to denote the corresponding graph.

### Model I

This model is the *average binding* gene-regulation model analysed in [19]. Considered more generally, the graph structure is the hypercube ℋ_*n*_ for *n* = 4, representing the binding of a single ligand to *n* sites, and the output is *fractional saturation*, or the average fraction of bound sites. Using the set theory notation described above, let |*S*| denote the number of elements in the subset *S*. The average binding input-output response is given by,

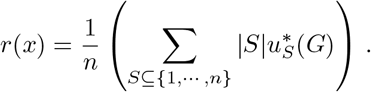

In [19], the ligand is a transcription factor (TF) binding to DNA and the output represents the normalised level of mRNA expression. The parameters are the independent association constants, *K*_*i,S*_ with *i < S*, introduced previously.

The Model I region forms a substantially reduced part of Ω_4_, although it still exhibits the cusp that approaches the (*p, s*) value of ℋ_4_ (Fig.S6).

### Model II

This model is the extended gene-regulation model analysed in [15]. The graph structure is the hypercube 𝒞_*n*+1_ for *n* = 4, representing the binding of one ligand, *A*, to *n* binding sites and the binding of a second ligand, *B*, to a single additional site (Fig.1A). In [15], *A* is a TF and *B* is RNA Polymerase being recruited to its promoter site.

It will be helpful for Models V and VI below to consider the more general situation of two ligands, *A* and *B*, with the input *A* binding to *a* sites 1, *· · ·, a* and the co-regulator *B* binding to *b* sites *a* + 1, *· · ·, a* + *b*. The graph structure is then the product 𝒞_*a*+*b*_ = 𝒞_*a*_ *×* 𝒞_*b*_ [16]. The vertices can be denoted by the ordered pair, (*S, T*), where *S* ⊆ {1, *· · ·, a*} and *T* ⊆ {*a* + 1, *· · ·, a* + *b*}. Independent association constants can be found for this model by following the spanning tree procedure described in [19],

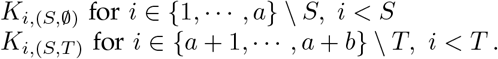

The first set of association constants are for *A* binding only in the presence of *A*, while the second set of association constants are for *B* binding in the presence of both *A* and *B*. All other association constants can be rationally expressed in terms of these independent ones [19].

For Model II, *a* = *n* = 4 and *b* = 1 and the output is the probability of *B* being bound when at least one site is bound by *A*. The input-output response for the resulting graph *G* is therefore given by,

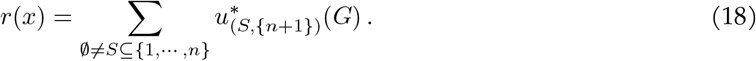

To fix the concentration of *B*, we chose *K*_*n*+1,(∅,∅)_[*B*] = 1.

Gene-regulation functions are sometimes found to increase monotonically with increasing TF concentration [15] and we were curious about the impact of such monotonicity on the (*p, s*) region. We therefore sought parametric conditions that would ensure this behaviour. We show below that if the affinity of *B* binding increases with the number of *A* sites that are bound, irrespective of which sites are bound, so that *K*_*n*+1,(*S*,∅)_ *> K*_*n*+1,(*T*,∅)_ whenever |*S*| *>* |*T* |, then *r*(*x*) increases monotonically with *x*. We imposed this constraint, which leads to the asymptotic (*p, s*) region shown for Model II in Fig.S6. Interestingly, the Model II region matches Ω_4_ better than does Model I, except when position is high and steepness is low.

**Figure S6:**
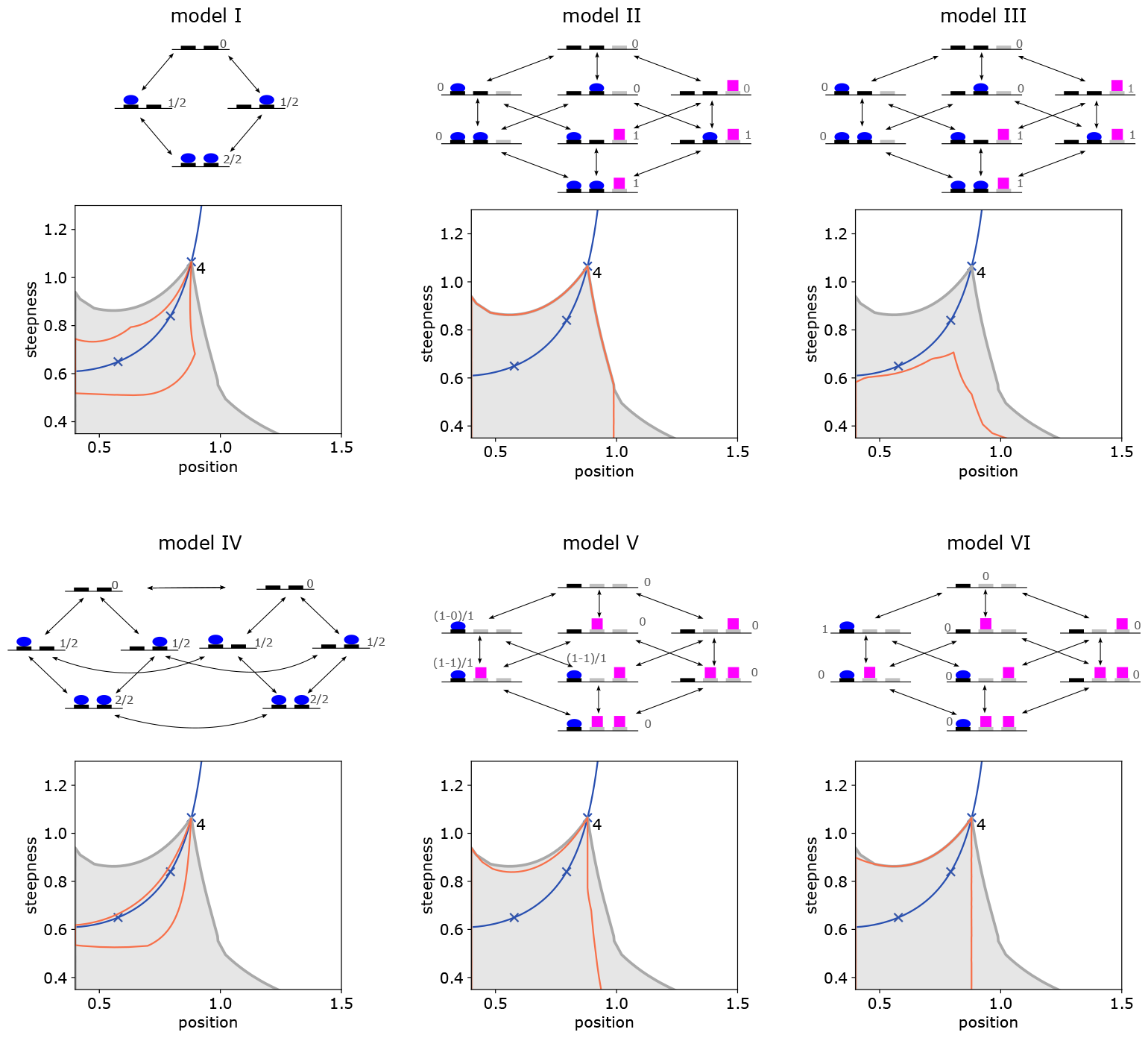
Asymptotic (*p, s*) regions for 6 specific models with *m* = 4 input binding sites. The red curve is the asymptotic boundary of each model, with the gray region showing the universal region Ω_4_. The same conventions are used as for Fig.3A in the main text, with the (*p, s*) value of ℋ_4_ marked at the cusp of Ω_4_. In each case the model region is contained within Ω_4_, as expected for universality. The graphs above each plot describe the corresponding model with the number of input binding sites taken to be 2 (Models I-IV) or 1 (Models V and VI), rather than 4, for ease of visualisation. The reversible edges are shown as bidirectional arrows and vertices are marked by expressions that give the corresponding weight in the input-output response.

### Model III

This model is a variation of Model II in which the monotonicity constraints on the parameters were dropped and the output was altered to allow for a response when no *A* is bound, so that,

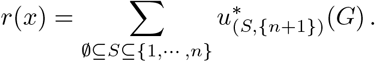

Fig. S6 shows that the asymptotic region is now quite different from that for Model II. The region has a poorly-defined cusp that is not on the Hill line and is far from the (*p, s*) value of ℋ_4_. This appears to be due to the choice of [*B*] being set by taking *K*_*n*+1,(∅,∅)_[*B*] = 1. If *K*_*n*+1,(∅,∅)_[*B*] is taken to be sufficiently large, the region acquires a well-defined cusp on the Hill line that approaches ℋ_4_, as in Fig.1C in the main text, which shows the (*p, s*) region for this model with *K*_*n*+1,(∅,∅)_[*B*] = 1000. Model III reveals the impact on the (*p, s*) region of apparently small changes in the input-output response and also the influence of other ligands, a topic that deserves further study.

### Model IV

This model is the Monod-Wyman-Changeaux model for allostery [23], as analysed using coarse graining in [24]. A biomolecule is assumed to exist in two inter-converting conformations, to which a ligand binds at *m* = 4 sites. We follow the original parameterisation, with *L* being the label ratio for inter-conversion of the conformations with no ligand bound, and *K*_1_ and *K*_2_ being the association constants for binding to each conformation independent of the site or the pattern of binding at other sites. The output is fractional saturation, as in Model I. The input-output response is given by the Monod-Wyman-Changeux formula [24, Eq.55],

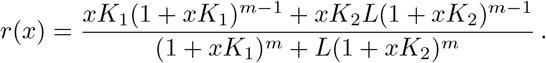

The asymptotic (*p, s*) region shows a clear cusp on the Hill line which approaches *H*_4_ (Fig.S6). The upper boundary of the region runs close to the Hill line and the lower part of the region is substantially restricted.

### Model V

This model considers the binding of an activator *A* to sites 1, *· · ·, a*, and the binding of a repressor *B* to sites *a*+1, *· · ·, a*+*b*, using the notation introduced for Model II above. Here, *a* = 4 and *b* = 2. Each vertex (*S, T*) is taken to influence the output by the amount ι (*S, T*) = max(0, |*S*| − |*T* |) and the input-output response is taken to be the fractional average of this quantity,

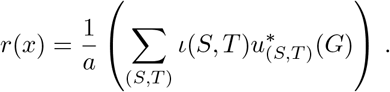

For this model, we took [*B*] = 1 in the same concentration units used for the association constants. The asymptotic (*p, s*) region for this model matches Ω_4_ less well than Model II and fails to extend to regions of higher position.

### Model VI

This model is a variant of Model V in which the repressor *B* is considered to be dominant over *A* and all *A* sites have to be occupied for output to occur. The graph is the same and the concentration of *B* was fixed in the same way but the input-output response is now taken to be,

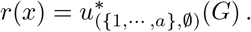

It is interesting that the asymptotic (*p, s*) region for this model is quite similar to that for Model V, despite the substantial difference in the output.

## 8 Monotonicity of the input-output response for Model II

In Model II above, we restricted the input-output response, *r*(*x*), to be a monotonically strictly increasing function of *x*, so that *r*(*x*) *< r*(*y*) whenever *x < y*. We prove here the parametric conditions that result in this behaviour. We require two simple observations.

### Lemma 1.

Suppose *a*_0_, *· · ·, a*_*n*_ and *b*_0_, *· · ·, b*_*n*_ are positive numbers. If *a*_*k*_*/b*_*k*_ does not decrease with *k*, so that *a*_*k*_*/b*_*k*_ ≤ *a*_*l*_*/b*_*l*_ when *k < l*, then the rational function

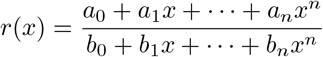

does not decrease for *x >* 0. If *a*_*k*_*/b*_*k*_ also increases at some *k*, so that *a*_*k*_*/b*_*k*_ *< a*_*k*+1_*/b*_*k*+1_, then *r*(*x*) increases strictly for *x >* 0.

**Proof**. The numerator of the derivative of *r*(*x*) has the form

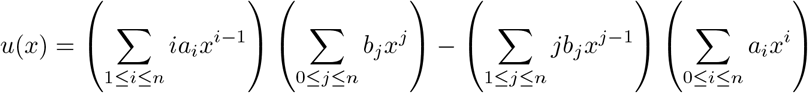

and the sign of *u*(*x*) determines the monotonicity of *r*(*x*). If *u*(*x*) ≥ 0 for *x >* 0, then *r*(*x*) is nondecreasing for *x >* 0 and if *u*(*x*) *>* 0 for *x >* 0, then *r*(*x*) is strictly increasing for *x >* 0. We may reorganise the expression for *u*(*x*) so that,

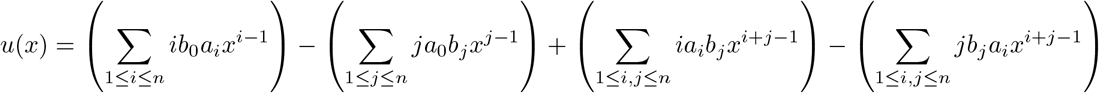

Collecting together the same powers of *x*, the first two terms yield the expression

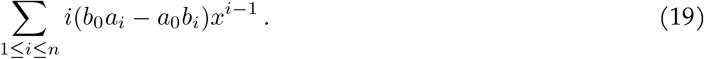

Doing the same for the last two terms, we see that the resulting expression can be broken into two parts,

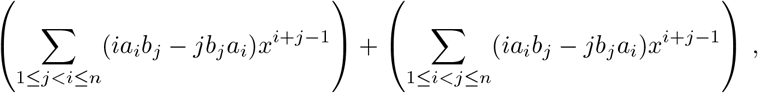

where the terms corresponding to *i* = *j* have cancelled. Since the indices *i* and *j* are arbitrary, we are at liberty to exchange them in the sum on the right and collect the two sums together to yield,

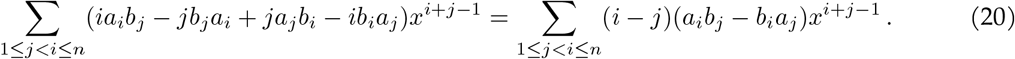

Because the *a*’s and *b*’s are all positive, it follows from Eqs.19 and 20 that, if *x >* 0 and *a*_*k*_*/b*_*k*_ does not decrease with *k*, then *u*(*x*) ≥ 0 and that if *a*_*k*_*/b*_*k*_ increases for some *k*, then *u*(*x*) *>* 0 for all *x >* 0. The result follows, as claimed.

There is a minor extension of Lemma 1 which will be helpful below. If the numerator polynomial in *r*(*x*) has no constant term, so that *a*_0_ = 0, but all the other *a*’s and *b*’s remain positive, then the expression in Eq.19 becomes positive. Moreover, at no stage in the remainder of the proof is it necessary to divide by *a*_0_. Hence, even if the weaker conditions of the Lemma hold, and *a*_*k*_*/b*_*k*_ does not decrease with *k*, the fact that *a*_0_ = 0 ensures that *r*(*x*) increases strictly for *x >* 0.

### Lemma 2.

Suppose *I* and *J* are arbitrary finite index sets and that *a*_*i*_, *b*_*i*_ and *A*_*j*_, *B*_*j*_ are arbitrary positive quantities indexed over *I* and *J*, respectively. If *a*_*i*_*/b*_*i*_ ≤ *A*_*j*_*/B*_*j*_ for any *i* ∈ *I* and any *j* ∈ *J*, then

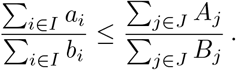

If the former inequality is strict for some *i* and *j*, then the latter inequality is also strict.

**Proof**. The inequality in the statement of Lemma 2 is equivalent to

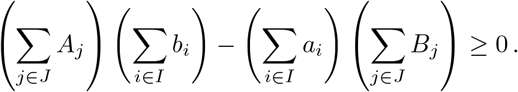

Rearranging the expression on the left gives,

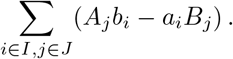

Since *a*_*i*_, *b*_*i*_, *A*_*j*_, *B*_*j*_ *>* 0, this expression is non-negative if *A*_*j*_*/B*_*j*_ ≥ *a*_*i*_*/b*_*i*_ for all *i* ∈ *I* and *j* ∈ *J*. Furthermore, if *A*_*j*_*/B*_*j*_ *> a*_*i*_*/b*_*i*_ for some *i, j*, the expression is positive. The result follows.

We can now return to the input-output response for Model II, as given in Eq.18. Recall from Eq.4 in the main text that s.s. probabilities are determined by the vector *µ*(*G*) at t.e. We can express the components *µ*_(*S,T*)_(*G*) in terms of the independent parameters introduced above for Model II as follows. Note that, for the case we are considering, in the notation used above, *a* = *n* and *b* = 1. Given any subset, *S* ⊆ {1, *· · ·, n*}, with |*S*| = *k*, suppose that the indices of *S* are written in increasing order, *S* = {*i*_1_, *· · ·, i*_*k*_} with 1 ≤ *i*_1_ *< i*_2_ *< · · · < i*_*k*_ ≤ *n*, and let *K*_*S*_ denote the product of independent association constants,

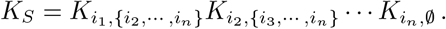

It then follows that,

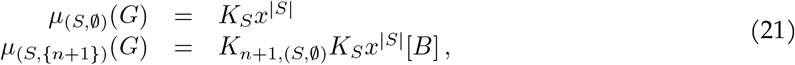

where *x* = [*A*]. This covers all the vertices of *G*. If we assemble the following two polynomials in *x*,

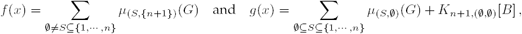

then Eq.4 of the main text shows that the input-output response in Eq.18 takes the form,

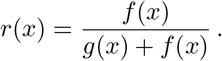

It is easy to see that *r*(*x*) is increasing if, and only if, *f* (*x*)*/g*(*x*) is increasing. This rational function falls under the scope of Lemma 1 and the minor extension discussed after the proof, in which the constant term in the numerator is 0. Accordingly, we must consider the ratio of the coefficients having the same power of *x*,

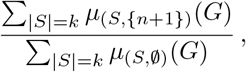

and show that these increase with *k* for 1 ≤ *k* ≤ *n*. This question now falls under the scope of Lemma 2. Choose 1 ≤ *k < l* ≤ *n* and apply Lemma 2 with the index set *I* being those subsets *S* with |*S*| = *k* and the index set *J* being those subsets *T* with |*T* | = *l*. Then, it follows from Eq.21 that,

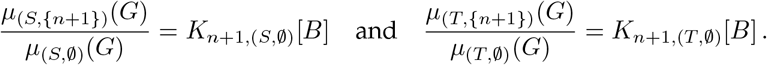

By hypothesis, *K*_*n*+1,(*S*,∅)_ *< K*_*n*+1,(*T*,∅)_ when |*S*| = *k < l* = |*T* |, so, applying Lemma 2, we can conclude that *r*(*x*) is monotonically strictly increasing, as claimed. This completes the proof of monotonicity for the input-output response of Model II.

## Notes

### Competing Interest Statement

The authors have declared no competing interest.

